# Genetic score omics regression and multi-trait meta-analysis detect widespread *cis*-regulatory effects shaping bovine complex traits

**DOI:** 10.1101/2022.07.13.499886

**Authors:** Ruidong Xiang, Lingzhao Fang, Shuli Liu, George E. Liu, Albert Tenesa, Yahui Gao, CattleGTEx Consortium, Brett A Mason, Amanda J. Chamberlain, Michael E. Goddard

## Abstract

To complete the genome-to-phenome map, transcriptome-wide association studies (TWAS) are performed to correlate genetically predicted gene expression with observed phenotypic measurements. However, the relatively small training population assayed with gene expression could limit the accuracy of TWAS. We propose Genetic Score Omics Regression (GSOR) correlating observed gene expression with genetically predicted phenotype, i.e., genetic score. The score, calculated using variants near genes with assayed expression, provides a powerful association test between *cis-*effects on gene expression and the trait. In simulated and real data, GSOR outperforms TWAS in detecting causal/informative genes. Applying GSOR to transcriptomes of 16 tissue (N∼5000) and 37 traits in ∼120,000 cattle, multi-trait meta-analyses of omics-associations (MTAO) found that, on average, each significant gene expression and splicing mediates *cis*-genetic effects on 8∼10 traits. Supported by Mendelian Randomisation, MTAO prioritised genes/splicing show increased evolutionary constraints. Many newly discovered genes/splicing regions underlie previously thought single-gene loci to influence multiple traits.

## Introduction

Genome-wide association studies (GWAS) test millions of genome variants, such as single nucleotide polymorphisms (SNPs), for association with quantitative traits. Significant associations map a quantitative trait locus (QTL) to a genomic region tracked by the associated variant. Since most associations involve non-coding genetic variants, gene regulation is expected to mediate the effects of many QTL. While this can be investigated by associating gene expression with complex traits, the two measurements are not always on the same individuals and this prevents the direct association analysis. However, it is possible to find a genetic association between predicted gene expression and complex traits where the prediction of gene expression is made from SNP genotypes in the individuals with complex trait phenotypes, and the prediction equation is trained in other individuals with SNP genotypes and gene expression measurements. This is commonly referred to as a transcriptome-wide association study (TWAS) ^1,2^.

The power of TWAS is determined by the accuracy of predicting gene expression from SNP genotypes and by the proportion of phenotypic variance explained by the expression of a single gene. Most datasets with gene expression measurements are not large and the prediction of gene expression is often limited to *cis* eQTL because *trans* effects are small and so hard to estimate accurately. Also, given the polygenicity of most complex traits, the predicted expression based on cis eQTL of a single gene is likely to explain only a small proportion of the variance of a complex trait.

Here we propose an alternative approach to estimating the genetic association between gene expression and complex traits which we call genetic score omics regression (GSOR). The datasets with complex trait measurements and SNP genotypes are often very large and can be used to train a prediction equation that predicts complex trait genetic values from SNP genotypes. The prediction is called an estimated breeding value (EBV) in animals and plants or a polygenic score (PGS) in humans ^3^. Then this prediction equation can be applied to individuals with actual gene expression measurements to correlate gene expression with EBV or PGS. Potentially, GSOR has two advantages over traditional TWAS. Firstly, the prediction of EBV/PGS should be more accurate because it is trained on a much larger data set than the prediction of gene expression. Secondly, the part of the EBV/PGS due to effects of SNPs close to the gene, i.e., the local EBV^4,5^/PGS, can be calculated and correlated with gene expression. These SNPs are also those responsible for *cis* eQTL, so the test for a correlation between *cis* effects on gene expression and complex trait EBV is more powerful than in TWAS. The above description of GSOR assumed the use of gene expression measurements, but it could be applied to any omics phenotype. Here we use gene expression together with RNA splicing.

GWAS are often followed by a meta-analysis of the effect (beta and se) of variants. Where GWAS summary statistics are available for several traits, a multi-trait meta-analysis of GWAS can be used to identify variants affecting multiple traits ^6^, i.e., pleiotropy. Similarly, TWAS or GSOR also produces association summary statistics between gene expression and multiple phenotypes. Therefore, a multi-trait meta-analysis can also be applied to such summary data to investigate the pleiotropic effects mediated by regulatory mechanisms.

Understanding causal mechanisms behind QTL is important but challenging. In humans, large-scale GWAS of both conventional and molecular phenotypes such as gene expression ^7^ and RNA splicing ^8^ improved the understanding of QTL causal effects. In animals, only a few causal QTL are identified and one of the most extraordinary QTL is a mutation in the gene for diacylglycerol O-acyltransferase 1 (*DGAT1*) in cattle. This single QTL explains 30%-40% of the phenotypic variance of milk production traits ^9,10^. While this QTL was previously identified to be caused by a protein-coding mutation ^9,11,12^, more recent studies indicated regulatory effects ^10,13^, possibly due to multiple causal mutations. The new CattleGTEx ^14^ and the FAANG consortium ^15^ provide opportunities to explore the causal regulatory mechanisms behind this QTL.

A complication with the interpretation of GWAS and TWAS results is caused by linkage disequilibrium (LD). One SNP may be associated with the expression of a gene and with a complex trait because of LD between this SNP and both a QTL for the trait and an eQTL for gene expression. However, if all SNPs that affect the expression of the gene have a proportional effect on the complex trait, then this is evidence that the gene expression causes variation in the complex trait. This is the logic of Mendelian randomization as implemented in SMR ^16^, which we use here to validate our results and explore causality. In addition, genes with important functions in mammals may have undergone purifying selection or are under evolutionary constraints across species. In this paper, we also investigate whether prioritised putatively causal genes show evidence of purifying selection.

We developed and applied GSOR to transcriptomes from 16 tissues from ∼5,000 cattle and 37 complex phenotypes from 113,000 cattle to dissect the genetic effects on complex traits mediated by the transcriptome. We propose a meta-analysis to quantify the pleiotropic effects of regulatory loci. We then use Mendelian Randomisation and the ratio of nonsynonymous substitution (dN) to synonymous substitution rates (dS) ^17^ to verify these effects and combine GSOR and SMR to dissect causal regulatory mechanisms. We show that blood group genes *ABO* and *ACHE* (Cartwright blood group) mediate causal effects on protein concentration and mastitis via expression and splicing, supporting conserved and widespread regulatory effects on mammalian complex traits.

## Results

### Genetic Score Omics Regression (GSOR)

It is more likely for a population with phenotypic records to have a larger sample size than a population with omics datasets, such as gene expression and RNA splicing. Therefore, we developed GSOR which estimates the effects (*b*) of omics features (Ω) on a complex phenotype leading to an EBV or PGS, *ĝ*_P_. Details are given in the Methods section, but the basic form of GSOR is:

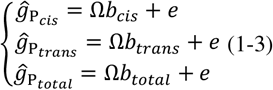

The response variable is the EBV or PGS for a complex trait (*ĝ*_P_) estimated using either genetic variants close (±1Mb of TSS) to a gene 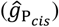 or all other genetic variants 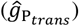, or is the total EBV/PGS 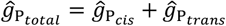. Accordingly, *b*_*cis*_, *b*_*trans*_ and *b*_*total*_ is the coefficient of regression of 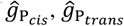 and 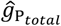, on the gene expression or splicing value. GSOR allows the fitting of random effects of a relationship matrix to control for population structures or confounding factors. GSOR is freely available at https://github.com/rxiangr/GSOR-and-MTAO.git.

To compare GSOR with the conventional TWAS, we analysed simulated and real data of 113,000 cattle with 16M sequence genotypes and 37 complex trait phenotypes and 945 cattle with 6M sequence genotypes and gene expression in blood (See Methods). To match with the implementation of GSOR, TWAS was also conducted using linear mixed models where gene expression predictors were trained by jointly fitting two genomic relationship matrices (GRMs) built using *cis* and *trans* variants. The predicted gene expression was then correlated with the complex trait phenotypes.

Using real bovine genotype data and ARS-UCD1.2 genome coordinates, we simulated causal *cis* and *trans* eQTL for 16,600 genes and created 5 scenarios of nulls where causal eQTL and causal QTL did not overlap and 5 alternative scenarios where causal eQTL and causal QTL did overlap (see Methods). Using Receiver Operating Characteristic (ROC) analysis of results, we showed that GSOR outperformed TWAS in detecting causal genes based on *cis*-predicted gene expression (Figure 1a,b) and *cis+trans* predicted gene expression (Supplementary Figure 1a,b). In the current simulation framework, neither GSOR nor TWAS had the power to detect causal genes based on *trans*-predicted gene expression (Supplementary Figure 1c,d).

**Figure 1.**
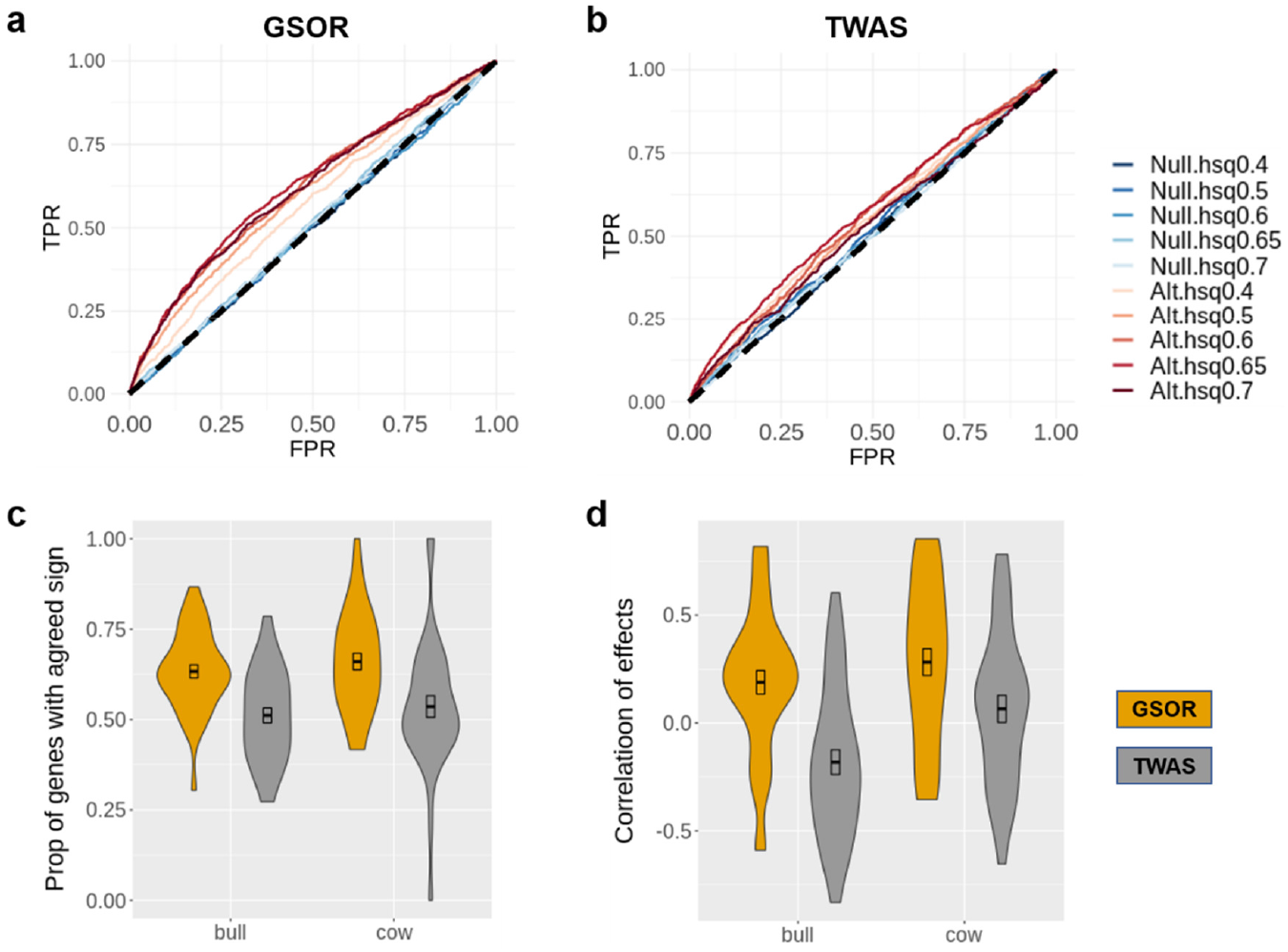
Comparison of results between GSOR and TWAS using simulations and real data. Receiver Operating Characteristic (ROC) analysis of results from GSOR and TWAS using simulated data are shown in (**a**) and (**b**), respectively. TPR: true positive rate. FPR: false positive rate. 10 scenarios were simulated with varying heritability (hsq) of traits. 5 traits were simulated under the null (Null) where no causal eQTL overlapped with causal QTL and another 5 traits were simulated under the alternative (Alt) scenarios where causal eQTL overlapped with causal QTL for more than 1000 genes. A comparison of results from real data is shown as violin plots in (**c**) and (**d**). In bull and cow datasets, the agreement of gene expression-phenotype association between *cis* and *trans* predicted values were compared. The comparison was based on the proportion of genes with the same sign (**c**) or correlation of effects (**d**), between *cis* and *trans* predicted values for 37 traits in each sex.

We next compared the results of GSOR and TWAS by analysing real blood gene expression data and 37 traits of 113,000 bulls and cows (Supplementary Table 1). If the increased expression of a gene causes a change in a complex trait, we expect the direction of that change to be the same for both *cis* and *trans* effects on gene expression. We observed that, in both sexes across 37 traits, the agreement of the direction of *cis* and *trans* effects and the correlation of effects between them was higher in GSOR than in TWAS (Figure 1c,d). In addition, we estimated the *π*_1_ value (lower bound on the proportion of truly alternative features ^18^, commonly used to indicate the proportion of replicated associations between different analyses ^7,14^) for results of GSOR and TWAS. We observed relatively higher *π*_1_ for GSOR than TWAS when replicating the gene expression-phenotype between *cis* and *trans* predicted *ĝ* (Supplementary Figure S2a), although for both GSOR and TWAS, *π*_1_ between *cis* and *trans* predicted *ĝ* was low. We also replicated the gene expression-phenotype association between bulls and cows, where we observed a much higher *π*_1_ for GSOR than TWAS using *cis* predicted *ĝ* (Supplementary Figure S2b). Overall, our results support the conclusion that GSOR has advantages over conventional TWAS in detecting genes whose expression is causally associated with complex traits.

### Multi-trait meta-analysis of omics-associations (MTAO)

We applied GSOR to transcriptome data (gene expression and RNA splicing) of 16 tissues combining newly generated data with data from CattleGTEx V0 ^14^ with a sample size > 100 (Supplementary Table 2) and *ĝ*_P_ of 37 cow traits (See Methods). Summary statistics of GSOR on gene expression and RNA splicing from 16 tissues across 37 traits of 103k cows are publicly available at https://figshare.com/s/c10ffab5abf329b1318f. To gain novel insights from vast summary statistics from GSOR, we introduce a multi-trait meta-analysis of omics-associations (MTAO) to quantify the extent of *cis* pleiotropy mediated by omics. MTAO estimates two statistics for each omics feature (including gene expression or splicing events) from each tissue: 1) the number of traits affected (*N*_*pleio*_) and 2) the magnitude of multi-trait effects (*M*_*pleio*_). The estimation of *N*_*pleio*_ adopted the method from Jordan 2019 ^19^ with increased rigor of significance testing (Methods). The estimation of *M*_*pleio*_ models the Chi-square distribution of signed t-values of each omic feature along with the correlation matrix of t-values across 37 traits to approximate the error covariance matrix (Methods). The R implementation of MTAO is publicly available at https://github.com/rxiangr/GSOR-and-MTAO/blob/main/README_MTAO.md. Note that MTAO can be applied to any results from omics-wide association testing including the conventional TWAS, as long as the regression coefficient (b) and standard error for each omics feature on the phenotype are obtained.

MTAO revealed that gene expression and RNA splicing mediate widespread *cis* pleiotropic effects on complex traits (Figure 2a-b). Across 16 tissues and 37 traits based on 3612 (SD=1037) significant genes (Supplementary Table 3), on average, the gene expression mediated *cis* pleiotropic effects on 9.7 traits (ranging from 8-13) with an average magnitude of 9.8 (ranging from 9-13). Based on 23,477 (SD=10633) significant introns (Supplementary Table 3), on average, splicing of an intron mediated *cis* pleiotropic effects on 8.4 traits (ranging from 8-11) with an average magnitude of 9.1 (ranging from 9-12) (Figure 2c-d).

**Figure 2.**
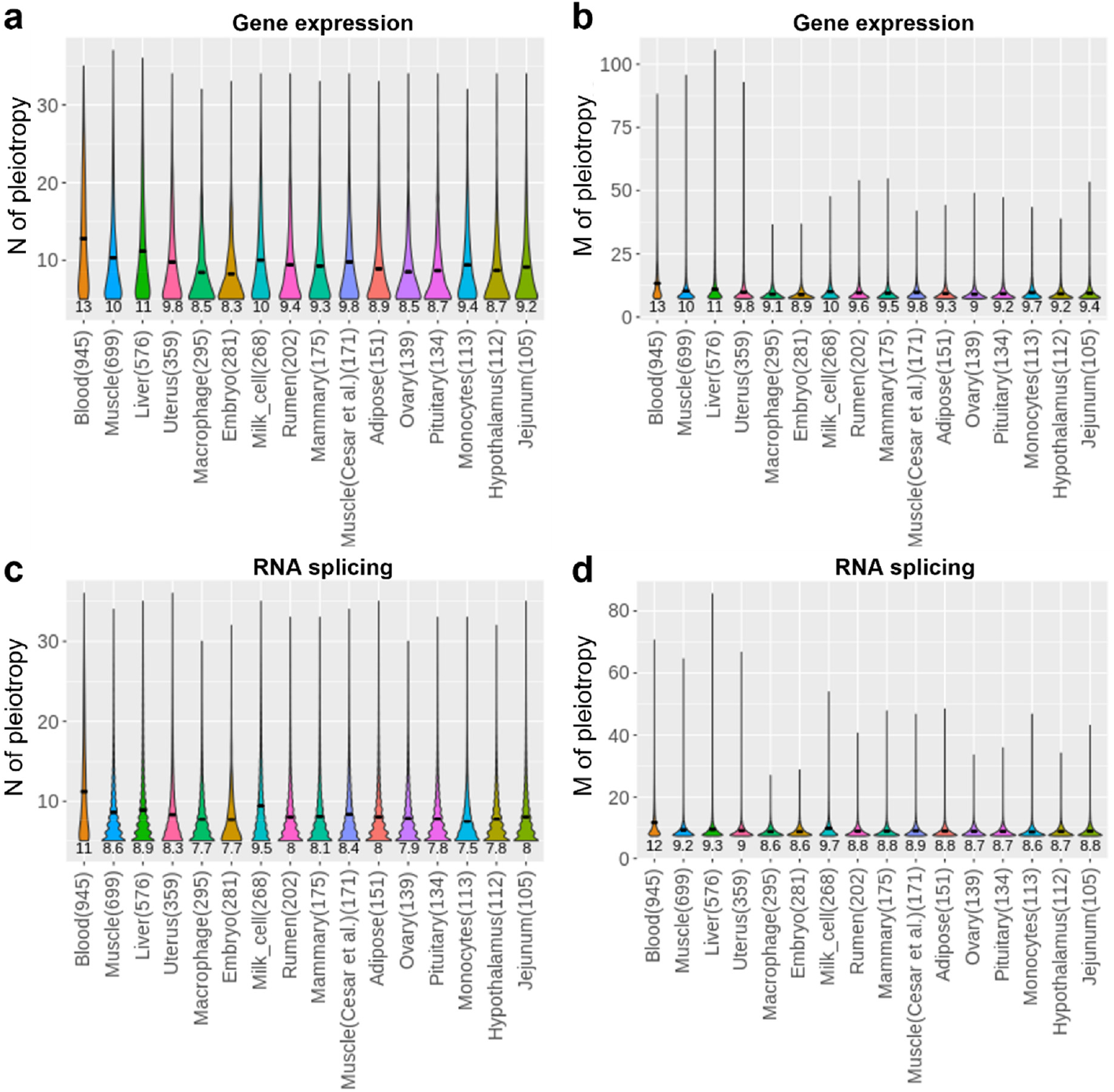
Gene expression and RNA splicing mediated *cis* pleiotropy. The average number of traits (N) associated with a gene expression (**a**) and the average magnitude (M) of pleiotropy with a gene expression (**b**); and the average N associated with a splicing event (**c**) and the average M of pleiotropy with a splicing event (**d**) are shown across 16 tissues. The numbers under each plot are the average values of N and M per tissue. The numbers in brackets are the sample size of each tissue.

### MTAO, Mendelian randomisation, and selection

MTAO identifies genes whose expression or splicing is associated with multiple traits. However, this association could be due to LD between eQTL or sQTL and QTL for complex traits. To test if variation in gene expression or splicing causes variation in complex traits, we conducted the summary data-based Mendelian randomization (SMR) in combination with the heterogeneity in dependent instruments (HEIDI) ^16^ test based on *cis* eQTL and sQTL mapped from 16 tissues and GWAS of 37 traits ^20,21^ (see Methods). HEIDI tests, including the more recent version using multiple top SNPs ^22^, for heterogeneity in the relationship between the effect of a variant on gene expression and the complex trait. Significant heterogeneity implies that the association between gene expression and trait is not causal and could be due to LD ^16,23^. Where results of SMR are available for multiple traits for a gene or an intron which passed the HEIDI test, we also used a multi-trait meta-analysis to combine SMR results across traits to identify genes or splicing events causing variation in more than one trait (See Methods). Then, we compared the genes/spliced introns prioritised by MTAO and by multi-trait SMR to check the extent of overlap (Figure 3a, Supplementary Table 4). Fisher’s exact tests show that the overlap of prioritised genes/spliced introns between MTAO and SMR is on average 2.4 times more than expected by random chance and is significant in most tissues (Figure 3a).

**Figure 3.**
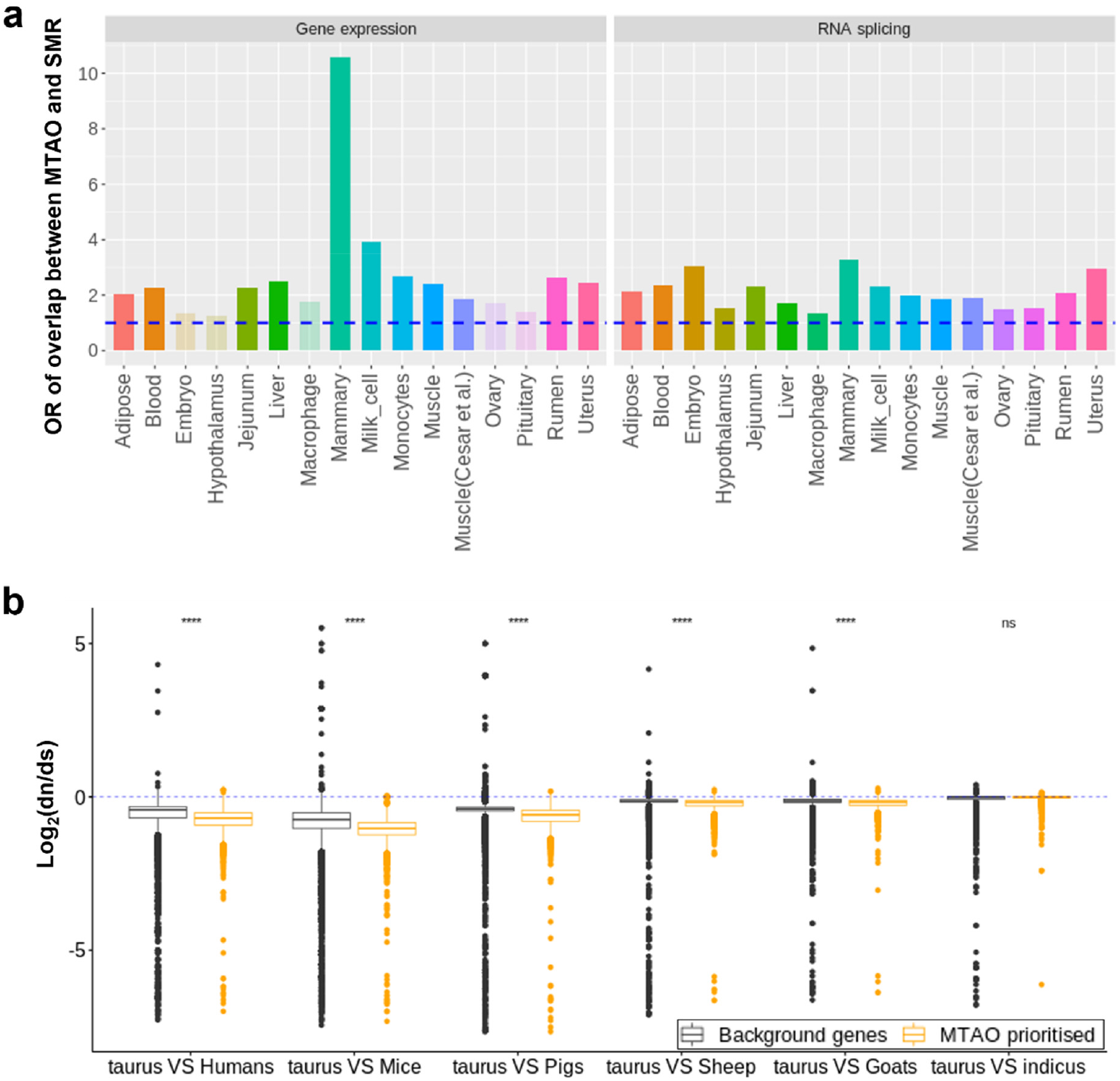
Supportive evidence for multi-trait meta-analysis of omics-associations (MTAO). **a**: Overlap of prioritised genes/introns between multi-trait meta-analysis of omics-associations (MTAO) and multi-trait summary data-based Mendelian randomization (SMR). Bars represent the odds ratio (OR) of fisher’s exact test of the overlap. The dashed blue line indicates OR = 1. Bars with transparent colors indicate the p-value of fisher’s exact test > 0.05 after multi-test adjustment (Embryo, Hypothalamus, Macrophage, Ovary and Pituitary in gene expression). **b**: nonsynonymous (dN) to synonymous substitution rate (dS) ratios for MTAO prioritised genes compared between *Bos taurus taurus* cattle (taurus) and other mammals (1-to-1 orthologs), including *Bos indicus* which is a sub-species of cattle. ^****^: t-test p-value < 1×10^−4^; ns: t-test not significant. In total 14504 ortholog genes participated in the analysis.

In addition, based on 1-to-1 orthology, we compared the dN/dS ratios of genes prioritised by MTAO between cattle and other species, including humans, mice and *Bos taurus indicus* which is a sub-species closely related to *Bos taurus taurus* cattle (Figure 3b). When comparing cattle and other species, MTAO prioritised genes showed significantly reduced dN/dS ratios than random genes. This suggests that MTAO prioritised genes show relatively stronger purifying selection ^17^ between cattle and other mammals, i.e., evolutionary constraint, and therefore that they play important functional roles in mammals.

### Detection of trait-relevant tissues

We next used results obtained from GSOR and SMR together to rank tissues according to their relationship with each trait. We used a heuristic index that combined results from GSOR, SMR, and HEIDI with the number of genes and individuals to rank tissues for each trait (see Methods) (Figure 4). Our analysis shows that blood, milk cells, liver and uterus had the highest and most consistent ranking in their connections with the traits analysed. This appears to be plausible given our collection of traits has a bias toward milk production and fertility. The ranking of tissues on average had a low correlation (spearman *rho* = 0.35) with the sample size and these correlations were not significant (Supplementary Table 5).

**Figure 4.**
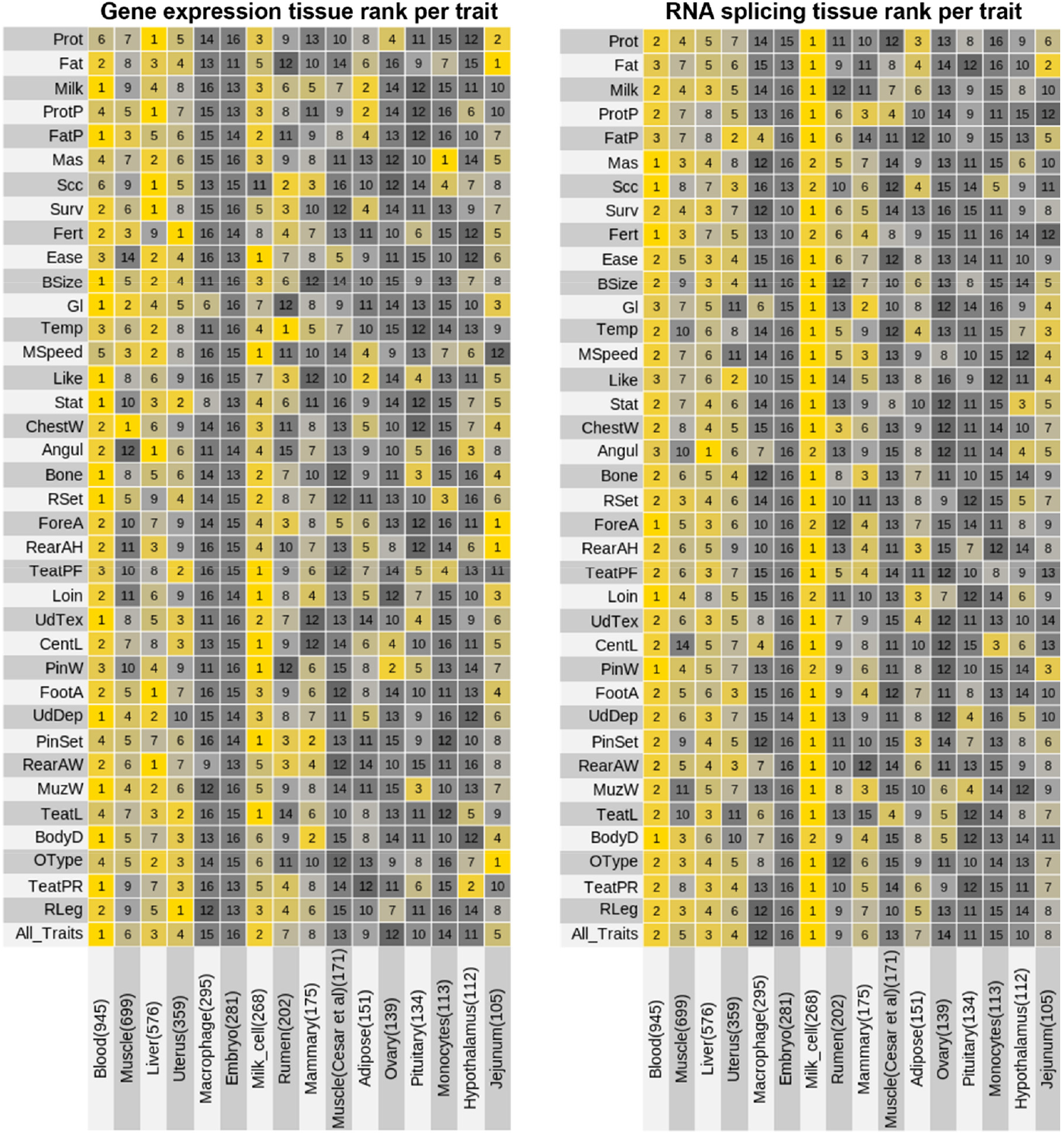
Ranking of tissues based on their importance to each trait. For each trait, we estimated the sum of the effects of GSOR and SMR across genes (or introns) per tissue adjusted by the number of genes and individuals. This sum was used to rank tissues for each trait. The numbers within cells indicate the tissue ranking from 1 to 16 per trait. The ranking for “All_Traits” is the ranking of tissues averaged across all traits. The numbers in brackets are the sample size of each tissue.

### Regulatory mechanisms underlie previously thought single-gene loci

The most understood yet controversial QTL in cattle is diacylglycerol O-acyltransferase 1 (*DGAT1*) previously identified to be caused by a protein-coding mutation and affects milk production traits ^9,11,12^. For the first time, we provide statistical evidence to support that gene expression and RNA splicing at *DGAT1* are causally linked to many traits in many tissues, and such causal links are not restricted to milk production traits (Figure 5a,b). Both MTAO and SMR found putative causal effects of *DGAT1* expression and/or splicing in blood, liver, mammary gland, milk cells, and uterus (Figure 5a-e). *DGAT1* expression and splicing had putative causal effects on milk production, mastitis (MAS, average correlation with milk production traits 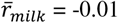), gestation length (Gl, 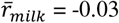), temperament (Temp, 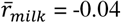), and stature (Stat, 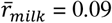). Therefore, our analysis supports the widespread regulatory causal effects of *DGAT1* on multiple traits possibly involving expression in many tissues. Another major QTL with regulatory effects reported in animals is *IGF2* (insulin-like growth factor 2) ^24^ and results show that RNA splicing of this gene was causally linked to many traits in tissues of liver and adipose (Supplementary Figure 3). Also, the expression of *MGST1* (Microsomal Glutathione S-Transferase 1)^25^ is causally linked to milk production traits in milk cells and the hypothalamus (Supplementary Figure 4). In addition, our current analysis did not observe causal regulatory effects from *GHR* (Growth Hormone Receptor, Supplementary Figure 5) with reported missense mutations ^26^ at sites conserved across vertebrates ^27^.

**Figure 5.**
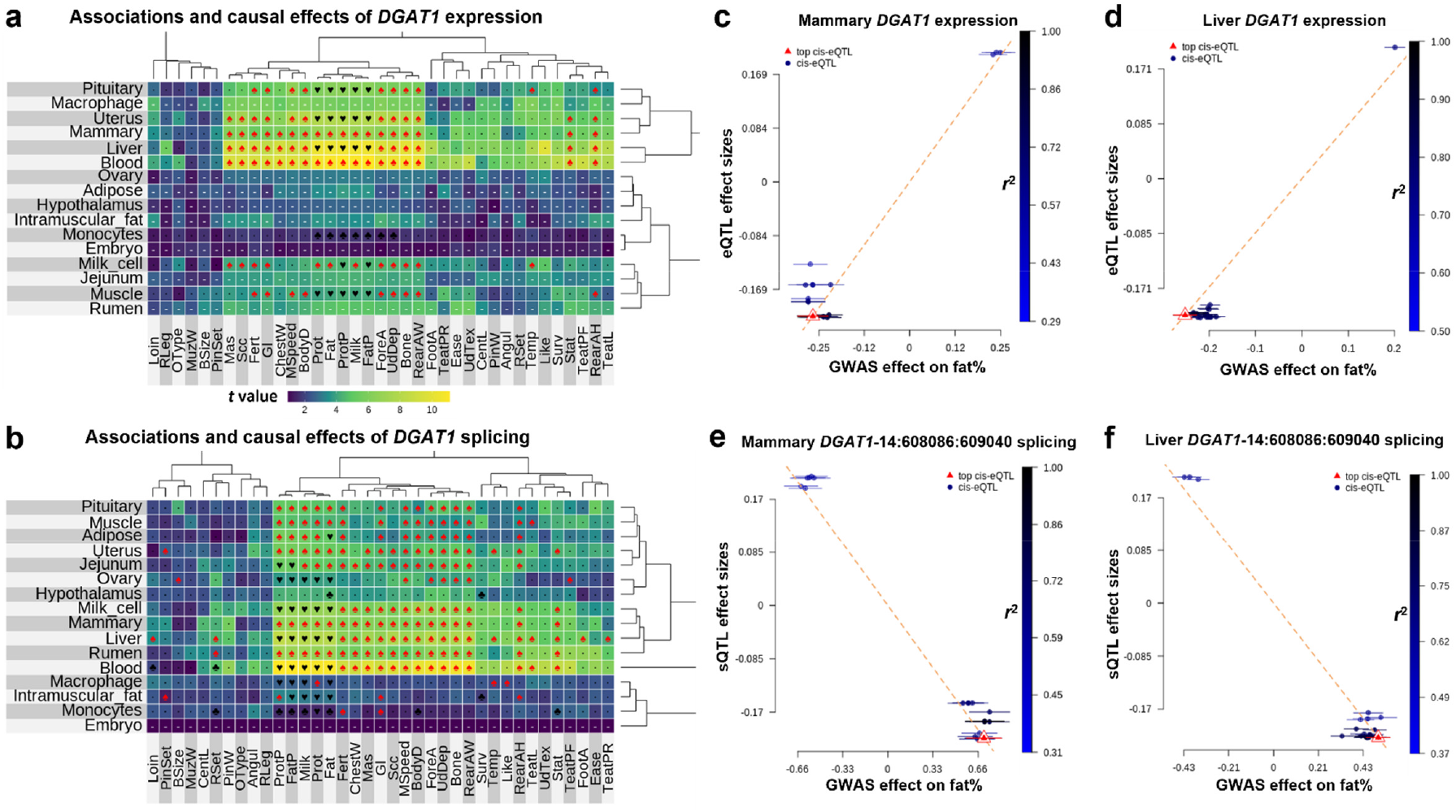
Widespread regulatory causal effects of *DGAT1*. The heat map of effects of *DGAT1* expression (**a**) and splicing (**b**) across tissues and traits based on GSOR. In these heat maps, red spades indicate causal effects inferred using summary-based Mendelian randomisation (SMR) independent of LD; black hearts indicate the causal effects confounded by LD while black clubs indicate causal effects without testing LD due to not enough SNPs. Black dots represent insignificant SMR test and white hyphens indicate no e/sQTL or QTL could be used for the SMR test. The dendrogram represents the hierarchical clustering of effects. The color scale of heatmaps is based on the magnitude of t (be/se) value of GSOR. **c**-**d** are examples of Mendelian randomisation using the expression eQTL and GWAS of fat%. **e**-**f** are examples of Mendelian randomisation using splicing sQTL and GWAS of fat%. 14:608086-609040 indicates the location of the spliced intron in *DGAT1*.

Based on the strongest statistical evidence from both GSOR and SMR for major traits of dairy cattle (non-linear assessment traits), we found that some traditionally recognised large-effect “single-gene” loci actually contain several adjacent genes or underlying spliced introns potentially causally linked to complex traits (Table 1 and Supplementary Data 1). We confirm the regulatory effects on milk production, gestation length, and height of several known causal loci, including *DGAT1* ^10^ via both gene expression and splicing, *MGST1*^25,28^ and *MATN3*^*29*^ via gene expression, and *CSF2RB*^*30,31*^ and *MUC1*^*32*^ via splicing. However, importantly, our evidence supports multiple regulatory loci underlying these major QTL. For example, there are 4 other genes (*ZNF34, IQANK1, LYNX1* and *SPAG1*) near *DGAT1* showing potentially causal effects on milk production traits independent of LD. Note that each of these genes used a different set of *cis* eQTL for SMR and HEIDI tests. For example, *ZNF34* and *IQANK1* are neighbor genes for *DGAT1*, and they had a relatively small number of shared *cis* eQTL with *DGAT1*, with an average LD-*r* = 0.7 with eQTLs of *DGAT1* (Supplementary Table 6). Even within *DGAT1*, there were 7 intronic regions whose splicing was potentially causally linked to cattle traits independent of LD (Supplementary Data 1).

**Table 1.**
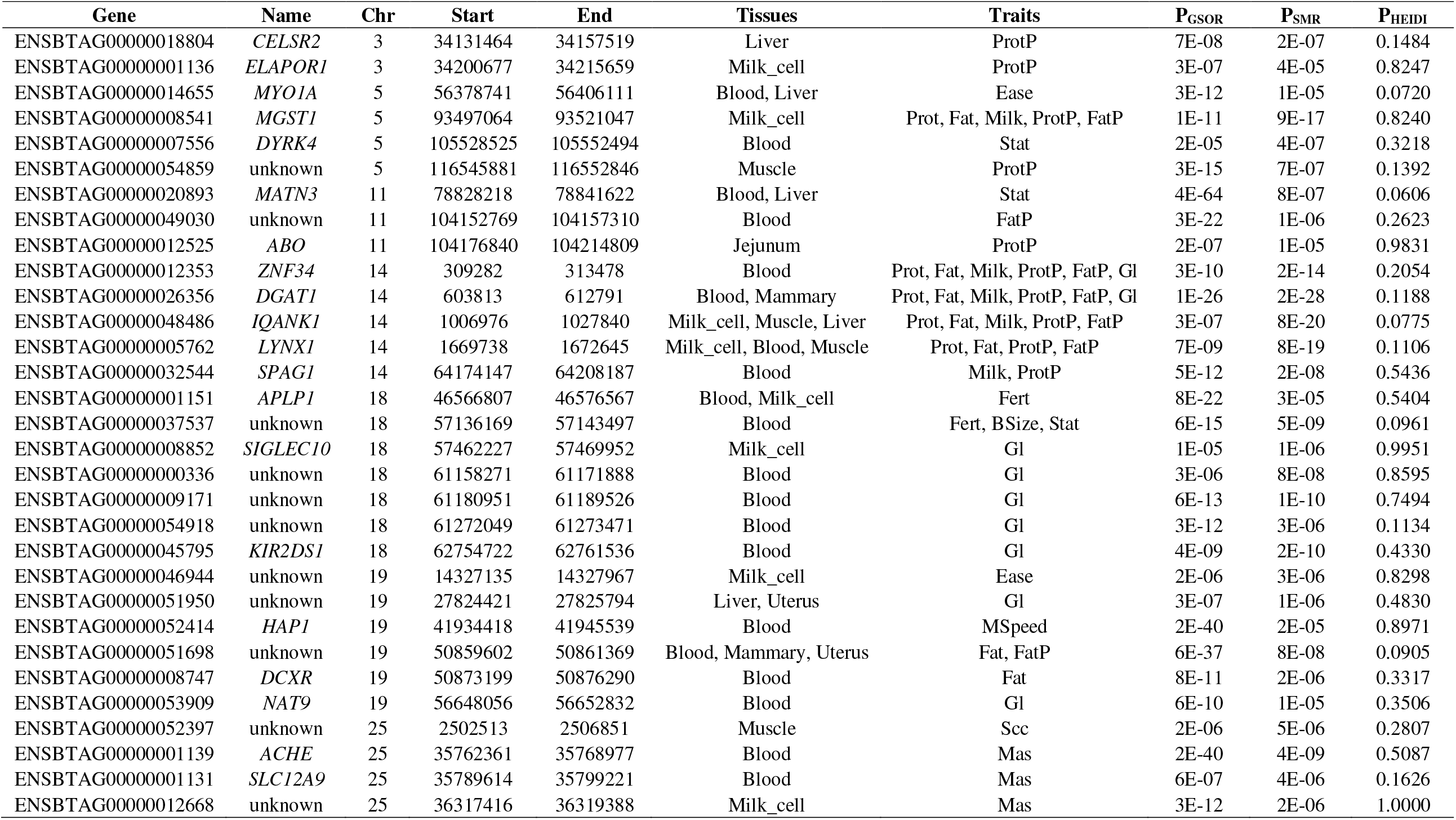
Summary of putatively causal links to cattle traits via gene regulation prioritised by GSOR (p-value shown in P_GSOR_) and SMR (p-value shown in P_SMR_ and P_HEIDI_). Note that P_HEIDI_ < 0.05 indicates LD confounding.

We identified many new or unannotated potentially causal loci, including two blood group genes, *ABO* (Histo-blood group) and *ACHE* (acetylcholinesterase, Cartwright blood group), causally linked to protein concentration and mastitis respectively via both gene expression and splicing in blood (Table 1 and Supplementary Data 1). Also, *DCXR* (dicarbonyl and L-xylulose reductase), which plays a significant role in glucose metabolism and causes human pentosuria (NCBI RefSeq) is linked to cattle fat yield via both gene expression and splicing in blood. *FLII* (FLII actin remodeling protein) causally linked to cattle gestation length via splicing in milk cells is a gene related to embryogenesis in Drosophila (NCBI RefSeq). Transcription factor gene *TAF9* (TATA-Box Binding Protein Associated Factor 9) is causally linked to cattle milk speed via splicing in blood.

## Discussion

The current study shows that GSOR has advantages over TWAS in finding genes whose expression or splicing is associated with complex traits. Many methods can be used to detect the association between gene expression and complex traits ^33-36^. However, the advantage of GSOR is due to the use of the large sample size to train the EBV or PGS, i.e., genetic score, for complex traits and the ability to calculate a part of this genetic score that is local to the gene whose expression is being considered leading to a more powerful test for *cis* eQTL or sQTL effects.

Applying GSOR and MTAO to the large dataset with transcriptomes and complex traits in cattle, we identified widespread genetic regulatory effects on complex traits, i.e., *cis* pleiotropic effects, mediated by the transcriptome. We show that on average each gene expression and splicing event mediate *cis* genetic effects on 10 and 8 traits, respectively, and this is comparable to our previous work where on average each variant affects 10 traits ^37^.

This suggests that the genetic effects mediated by *cis*-regulatory mechanisms on complex traits are prevalent. Using a multi-trait meta-analysis of SMR ^16^, a different approach to detect putative causal relationships between omics features and traits, we validate the results of MTAO. Also, we found that MTAO prioritised genes show significantly stronger purifying selection than random genes, supporting that these genes have important functions in cattle and other mammals. These pieces of evidence support that MTAO can be used to identify omics features, i.e., regulatory elements, of high importance to complex traits. Apart from gene expression and splicing, there are many other types of quantitative omics features such as the height of ChIP-seq peaks ^38^ and allele-specific imbalance ^27^ which can also be analysed by MTAO. Moreover, MTAO is summary-data based so it can be applied to any results from GSOR or conventional TWAS, as long as there are beta and standard error estimates for the association between the omics feature and the phenotype. Therefore, we expect MTAO to have an important place in future large-scale meta-analyses of different GSOR or TWAS studies in mammalian species.

We combined the results from the GSOR and SMR to prioritise tissues related to different phenotypes. Across different traits, blood, milk cells, liver, and uterus were the most informative tissues for traits analysed. Blood is one of the most common tissues/cell types sampled for omics studies due to its easy access. Milk cells can also be relatively easily accessed in dairy cattle ^39^. Although we have adjusted the analysis according to the sample size of each tissue, the tissue prioritisation might still have some biases towards those with larger sample sizes.

Understanding the causal nature of large-effect QTL provides opportunities for treatments. Here we used results from MTAO and SMR to dissect regulatory causal effects of some major cattle loci, including *DGAT1*. A protein-coding mutation in *DGAT1* was previously identified as the cause of this QTL’s effect on milk production traits, but there has been speculation that there are multiple causal variants in this region of the genome ^12,13^. Our results, for the first time, show a correlation between *DGAT1* expression and splicing and numerous traits. *DGAT1* expression in blood, liver, mammary gland, pituitary, milk cells, and rumen were correlated with milk and non-milk traits. This agrees with the pleiotropy model which we previously proposed ^32^ which is that QTL with a large effect on one trait is likely to have small effects on other uncorrelated traits.

In addition to *DGAT1*, the expression and splicing of 4 other genes (*ZNF34, IQANK1, LYNX1* and *SPAG1)* close to *DGAT1* was correlated with complex traits. Similar results were observed for other loci like *MGST1, MUC1*, and *CSF2RB*. This could be because mutations could affect the regulation of more than one nearby gene or it could be that these genes affect dairy traits directly. It has been demonstrated that multiple causal variants underlie human QTL ^40^. This may be the same for many traits of cattle, although we will need further experimental approaches to validate this.

We also provide statistical evidence for several new loci potentially affecting cattle traits via gene expression regulation, including blood group genes (*ABO* and *ACHE*). Interestingly, a deletion in *ABO* has been causally linked to pig complex traits via regulation of gut microbes ^41^ and here we found the regulatory effects of *ABO* in the jejunum (Table 1). However, there have not been reports regarding potentially causal effects via expression and splicing on complex traits linked to *ABO* or *ACHE* (Cartwright blood group). To our knowledge, it’s the first study to find *DCXR* related to glucose metabolism and *FLII* related to prenatal development, to have regulatory QTL in cattle.

In conclusion, we have introduced new methods and meta-analysis strategies to link omics information and complex traits. These methods are supported by the analysis of simulated and real data and by established Mendelian randomisation methods which account for LD. Our methods detected widespread pleiotropic effects mediated by multiple regulatory mechanisms and prioritised many genes and splicing events with potential causal associations with cattle traits. Primarily developed in cattle, GSOR and summary-data-based MTAO can prioritise informative omics-phenotype associations in any species.

## Methods

### RNA-seq data

The RNA-seq and genotype data analysed included those generated by Agriculture Victoria Research (AVR) in Victoria, Australia, and those provided by the CattleGTEx consortium ^14^ (Supplementary Table 2). The animal ethics was approved by the DJPR Animal Ethics Committee (application numbers 2013-14 and 2018-2019), Australia. Blood samples were taken from 390 lactating cows from 2 breeds, and milk samples from 281 lactating cows from 2 breeds. The processing of samples, RNA extractions, and library preparation followed that previously described ^28,42^. RNA sequencing (RNA-seq) was performed on a HiSeq3000 (Illumina Inc) or NovaSeq6000 (Illumina Inc) genome analyzer in a paired-end, 150-cycle run. Only RNA-seq data of 356 Holstein and 26 Jersey with > 50 million reads for milk cells or > 25 million reads for white blood cells and had concordant alignment rate ^43^ > 80% were used. QualityTrim (https://bitbucket.org/arobinson/qualitytrim) was used to trim and filter poor-quality bases and sequence reads. Adaptor sequences and bases with a quality score of <20 were removed. Reads with a mean quality score less than 20, greater than 3 N, greater than three consecutive bases with a quality score less than 15, or a final length of fewer than 50 bases were discarded. High-quality raw reads were aligned to the ARS-UCD1.2 bovine genome ^44^ with STAR ^43^ using the 2-pass method. The gene counts were extracted by FeatureCount ^45^. Leafcutter ^33^ was used to generate junction files which were then used to create the RNA splicing phenotype matrix, i.e., intron excision ratio ^33^.

The RNA-seq gene counts of 15 tissues (Supplementary Table 2) where the sample size > 100 were downloaded from CattleGTEx website http://cgtex.roslin.ed.ac.uk/. The blood counts generated by AVR (white blood cells) and CattleGTEx were combined. All gene counts were normalised by voom ^46^ and then underwent quantile normalisation for the following analyses. Junction files from CattleGTEx tissues were also downloaded and data from each tissue was processed with leafcutter ^33^ to generate RNA splicing phenotype. Milk cell data used in this study was only from AVR.

### Genotype data

The genotype data for Australian animals including those used for e/sQTL mapping (blood and milk cells) and association analysis of phenotypes (described later) consisted of 16,251,453 sequence variants imputed using Run7 of the 1000 Bull Genomes Project ^47,48^. The details of the imputation were described previously ^49^. Briefly, the imputation of bi-allelic sequence variants was performed with Minimac3 ^50,51^ and those variants with imputation accuracy *R*^*2*^ > 0.4 and minor allele frequency (MAF) > 0.005 in both bulls and cows were kept. Bulls were genotyped with either a medium-density SNP array (50K: BovineSNP50 Beadchip, Illumina Inc) or a high-density SNP array (HD: BovineHD BeadChip, Illumina Inc) and cows were genotyped with the BovineSNP50 Beadchip (Illumina Inc). The genotype data for CattleGTEx animals were generated previously ^14^ and included a total of more than 6 million sequence variants imputed also using Run7 of the 1000 Bull Genomes Project. Those variants with the imputation dosage R-squared > 0.8 and MAF > 0.001 were kept.

### Phenotype data

Data were collected by farmers and processed by DataGene Australia (http://www.datagene.com.au/) for the official May 2020 release of National breeding values. No live animal experimentation was required. DataGene provided the bull and cow phenotypes as de-regressed breeding values or trait deviations for cows, and daughter trait deviations for bulls (i.e., progeny test data for bulls). DataGene corrected the phenotypes for herd, year, season, and lactation following the procedures used for routine genetic evaluations in Australian dairy cattle. Phenotype data included a total of 8,949 bulls and 103,350 cows, including Holstein (6,886♂ / 87,003♀), Jersey (1562♂ / 13,353♀), cross-breed (36♂ / 5,037♀) and Australian Red (265♂ / 3,379♀) dairy breeds. In total, 37 traits were studied that related to milk production, mastitis, fertility, temperament, and body conformation and the details of these traits can be found in ^49^. For AVR blood samples, breed and days in milk (DIM) were fitted as fixed effects in the gene expression and splicing GWAS model. For the milk samples, experiment, DIM, and the first and second principal components, extracted from the expression count matrix, were fitted as fixed effects. Principle components were fitted to adjust for the high expression of the major milk protein genes, i.e., casein, in milk cells based on previous experiences ^28^.

### Genetic Score Omics Regression (GSOR)

A key feature of GSOR is the use of predicted phenotype value, i.e, genetic score (also called estimated breeding value or polygenic score), from a large reference population, as the explanatory variable to be associated with gene expression levels, splicing events, or other omic features. Another key feature of GSOR was the use of variants close to the gene whose expression is being studied to calculate a local or *cis* EBV/PGS. This would then be correlated with the expression or splicing of the gene. Note that although the local EBV/PGS was based on effects of SNPs near the gene, all SNP effects are trained jointly (described below). Where the total EBV/PGS minus the cis EBV/PGS was the trans EBV/PGS. It is generally recommended to use trait variant prediction models that jointly fit all variants together, such as gBLUP ^52,53^ or BayesR ^54,55^. Here we considered gBLUP for computational efficiency. A basic gBLUP model can be described as:

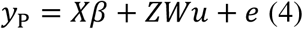

or

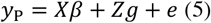

Where is *X* a design matrix, *y*_P_ is an n × 1 vector of phenotypes and n is the number of individuals; *β* is a vector of fixed effects and Z is the matrix allocating records to individuals; u is a vector of SNP effects with 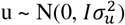 where I is an n × n identity matrix; *W* is a standardized genotype matrix and if models like GCTA ^56^ were used, 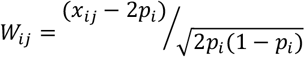 where *x*_*ij*_ is the number of copies of the 1^st^ allele for the i^th^ SNP of the j^th^ individual and *p*_*i*_ is the frequency of the 1^st^ allele; *g* is an n × 1 vector of the total genetic effects of the individuals with 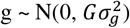 where *G* is the genomic relationship matrix (GRM) between individuals, 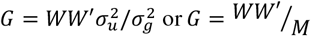 where 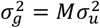 and M is the number of variants to explain 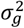; and e is the residual where 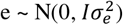. As Equation (4) and (5) are equivalent ^53,56,57^, it is possible to transform the BLUP of individual genetic score *g* to BLUP of *û*, i.e., SNP effects jointly estimated:

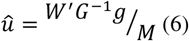

Equation 6 was implemented with GCTA BLUP ^56^ in the current study. Estimated *û* can be used to predict the genetic score, i.e., breeding value or polygenic score, of new individuals based on their genome-wide variant data: *ĝ* = *W*_*new*_ *û*. Because the SNP effects *û* was jointly estimated, it is also possible to use a subset of variants to predict *ĝ* (local EBV/PGS). For example, we have previously estimated *ĝ* of every 50kb windows of variants ^31^. In the current study, we estimate *ĝ* using variants close or distant to omic features such as genes or introns:

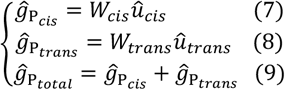

Where 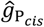 is the estimated genetic score using effects on phenotype (*û*_*cis*_) and the genotype matrix (*W*_*cis*_) of *cis* variants of omic features; 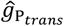 is the estimated genetic score using effects on phenotype (*û*_*trans*_) and the genotype matrix (*W*_*trans*_) of *trans* variants; and 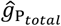 is the total genetic score by summing the *cis* and *trans* estimations. In the GSOR, for each gene, the *cis* variants were defined as ±1Mb of the transcription start site of the gene and the *trans* variants are the remaining variants. For each intron, *cis* variants were defined as those within 1Mb down and upstream of the intron (from intron start – 1Mb to intron end + 1Mb) and the *trans* variants are the remaining variants. Once *cis* and/or *trans* estimated genetic scores of genes/introns were obtained, they were analysed as response variables with gene expression or RNA splicing (intron excision ratio) as predictors:

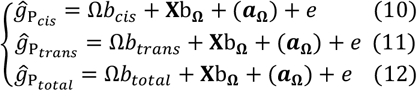

Where Ω is an n × 1 vector of omics values such as gene expression or RNA splicing corrected for other fixed effects such as breed, sex and experiments, *b*_*cis*_ is the regression coefficient of the *cis* estimated genetic score *ĝ*_*cis*_ *on* Ω, *b*_*trans*_ is the regression coefficient of the trans estimated genetic score *ĝ*_*trans*_ on Ω, and *b*_*total*_ is the regression coefficient of the total genetic score *ĝ*_*total*_ *on* Ω; **X** was the design matrix for fixed effects for data with omics measurements, e.g., breeds; *b*_**Ω**_ was the vector of fixed effects in the omics data; ***a***_**Ω**_ was a vector of random polygenic effects ∼N(0, **G**σ_g_^2^) which can be optionally fit to adjust confounding factors, **G** = genomic relatedness matrix (GRM) based on all variants and σ_g_^2^ = random polygenic variance, and e is the residual.

The implementation of GSOR was undertaken in R (v4.0.0) and is publicly available at (https://github.com/rxiangr/GSOR-and-MTAO). GSOR can work with or without random effects and when it does, it uses the implementation of eigendecomposition of the relationship matrix to speed up the variance components analysis. In the AVR high-performance cluster (slurm) system with 1 node, GSOR used 1.6G RAM and took 4.5 minutes to associate expression levels of 16,564 genes with the trait genetic score of 945 individuals fitting a GRM for each regression.

### Conventional Transcriptome-Wide Association Studies (TWAS)

Opposite to GSOR, a conventional TWAS essentially associates predicted gene expression in a large population with phenotypes of this population. The variant predictor was directly trained in the population where omics data was available. To make results from GSOR and TWAS comparable, we conducted TWAS using linear mixed model approaches and the variant predictors were trained using the omics data from blood which had the largest sample size across all tissues analysed. To train the variant predictor, a 2-GRM model was analysed for each omic feature which is similar to equation 5:

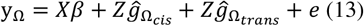

Where *y*_Ω_ is an n × 1 vector of omics values such as gene expression or RNA splicing, *β* is a vector of fixed effects; 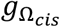 is an n × 1 vector of the total genetic effects of the individuals with 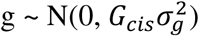 where *G*_*cis*_ is the GRM built by *cis* variants of the omic feature; 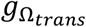 is an n × 1 vector of the total genetic effects of the individuals with 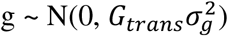) where *G*_*trans*_ is the GRM built by *trans* variants of the omic feature; Z is the matrix allocating records to individuals; e is the error term. Once 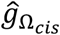 and 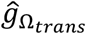 were obtained, equation (6) was applied to estimate SNP BLUP for omics data: 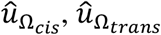 and 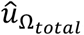, which were used to predict the omics scores, 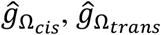 and 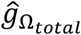 in the population with phenotypic records with equations 7-9. Then, predicted gene expression values were analysed as explanatory variables to associate with phenotypes:

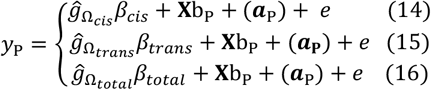

Where *y*_P_ is an n × 1 vector of phenotypes, *β*_*cis*_ is the regression coefficient for *cis* estimated omics score 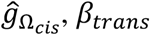 is the regression coefficient for *trans* estimated omics score 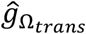, and *β*_*total*_ is the regression coefficient for the total omics score *ĝ*_*total*_; **X** was the design matrix for fixed effects, e.g., breeds; *b*_P_ was the vector of fixed effects in the dataset with phenotypic records; ***a***_P_ is random effects based on the genomic relationships between individuals with phenotypic data which can be optionally fit to adjust confounding factors and e is the residual. The training of variant predictors of omics data used gBLUP implementation of MTG2 and the TWAS used the implementation of OSCA ^58^.

### Simulations

To compare GSOR with TWAS, we simulated data where causal variants that affect gene expression and phenotypes were overlapped. We used the 6 million sequence genotypes from the blood dataset to simulate 16,600 gene expression phenotypes with the following framework: 1) gene coordinates from bovine ARS-UCD1.2 reference genome were used; 2) the expression of each gene had 1-2 causal cis eQTL and 0-3 causal trans eQTL (on different chromosome to the gene) so that all genes had causal cis eQTL but not all genes had causal trans eQTLand genes had causal cis eQTL only or both causal cis and trans eQTL; 3) across 16,600 genes, 1049 had causal cis and/or trans eQTLs overlapping with causal QTL under the alternative scenario (described later) 268 of which had causal trans eQTL overlapping with causal QTL; 4) in total, 1,771 causal eQTL in the expression data were also causal QTL; 5) the effects of cis causal eQTL were randomly sampled from a uniform distribution where the minimum was 0.05 and the maximum was 0.5; 6) the effects of trans causal eQTL were randomly sampled from a uniform distribution where the minimum was 1e-6 and maximum was 0.05; this was to make average effects of trans eQTL 10 times smaller than cis eQTL; 7) the heritability of the expression of genes was sampled from a normal distribution where the mean *h*^2^ was 0.25 and the standard deviation was 0.2; only positive values were allowed.

Sixteen million sequence genotypes from more than 100K cows were used to simulate cow phenotypes with the following framework: 1) 5000 causal variants were defined and used to simulate 10 traits; 2) the first 5 traits (1-5) were simulated under the null where their 5000 causal variants did not overlap with causal eQTLs. These 5 traits had heritabilities of 0.4, 0.5, 0.6, 0.65 and 0.7, respectively; 3) the second set of 5 traits (6-10) were simulated under the alternative scenario, with the same heritability settings, but the 5000 causal SNPs overlapped with causal eQTL as described above. All simulations used the framework from GCTA GWAS model ^56^.

### Comparison between GSOR and TWAS

In simulations, a gene was defined as a causal gene if it had both causal eQTL and QTL, i.e, the same SNP was both eQTL and QTL. All genes analysed were then classified as causal or non-causal and this was analysed using the Receiver Operating Characteristic (ROC) curves against p-values from GSOR and TWAS. ROC analysis used the R package ‘pROC’ and resultant ROC curves were presented using ggplot2. In analysing real data, we compared gene-trait associations between cis and trans-predictions. In GSOR, both cis and trans variants were used to predict genetic scores to be associated with gene expression. In TWAS, both cis and trans variants were also used to predict omics values to be associated with phenotypes. For a gene, while we could not expect its association with a trait to be significant based on both cis and trans predictions, we could expect its trait association to have the same direction of effect in both cis and trans predictions. Therefore, we compared the proportion of significant genes with the same direction of effect in both cis and trans predictions between GSOR and TWAS. Genes with p < 0.05 in both cis 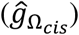 and trans predicted 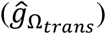 analysis in GSOR or TWAS were used for the comparison.

### Omics mediated cis pleiotropy

GSOR estimates the effects (beta) and standard error of each gene expression or splicing event on a trait. Combining these results across traits could provide insights into pleiotropy. Focusing on the cis-predicted genetic score, we performed a meta-analysis to quantify the extent of multi-trait effects of each gene expression or splicing event. The results of the analysis indicated the extent of cis pleiotropy mediated by omics. For each gene expression or splicing event, the t value (beta/se) from GSOR for each associated trait was obtained to model the number of traits (*N*_*pleio*_) affected and the magnitude (*M*_*pleio*_) of such pleiotropic effects. To estimate *N*_*pleio*_, the t values across traits were decorrelated using the Mahalanobis transformation described by Jordan et al. 2019 ^19^. Then, we adopted the method from Jordan et al 2019 ^19^ with a more stringent significance test:

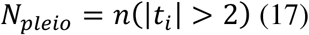

Where *n*(|*t*_*i*_| > 2) is the number of t values, out of the total number of K traits, of the omic feature *i* with a magnitude greater than 2. Two is used because it represents a standard t value in a normal distribution with a significance cutoff of p = 0.045. The significance test of *N*_*pleio*_ used 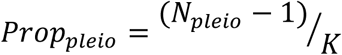 where *Prop*_*pleio*_ is the proportion of traits significantly affected by the omics feature. To obtain the p-value, *Prop*_*pleio*_ was then tested against the probability of 0.045 which is the probability of the t value being greater than 2 in the normal distribution. The reason for using 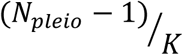 instead of using 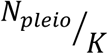 (used by Jordan et al. 2019^19^) is when *N*_*pleio*_= 1, i.e., the omics feature only affects one trait, then this does not qualify as pleiotropy, which is defined as genetic effects on more than 1 trait.

To estimate the magnitude of mediated pleiotropy, we used:

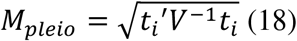

Where *t*_*i*_ is the effects (beta/se) from GSOR for each omics feature, *t*_*i*_′ is the transpose of *t*_*i*_ and *V*^−1^, *V* is the K ×K correlation matrix based on the t values for each trait. To obtain the significance of *M*_*pleio*_, 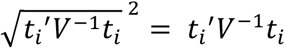 was tested against the *χ*^2^ distribution with degrees of freedom K. This approach was adopted from Bolormaa 2014 et al. ^6^ and Xiang et al. 2017 and 2020 ^32,59^ and gives identical results using the method from Jordan et al 2019^19^, but without the need for decorrelation of t values. The Rscript to conduct the meta-analysis of *N*_*pleio*_ and *M*_*pleio*_ are publicly available at https://figshare.com/s/c10ffab5abf329b1318f.

### Summary data-based Mendelian Randomization (SMR)

To verify the results from MTAO, we conducted SMR using mapped cis eQTL and sQTL (±1Mb from the gene or intron) from 16 tissues and GWAS results from 37 traits ^20^. Because cis eQTL or sQTL rely on SNPs very close to each other which usually have high LD, the heterogeneity in dependent instruments (HEIDI) ^16^ test is an effective analysis to distinguish causal from LD. The mapping of eQTL and sQTL are detailed in ^21^. Briefly, we first used a linear mixed model approach to map cis eQTL and sQTL in GCTA: *y*_Ω_ = *Xβ* + *Zg*_*all*_ *+ Wv + e* (19); where *y*_Ω_ is an n × 1 vector of omics values such as gene expression or RNA splicing, *β* is a vector of fixed effects like breeds, different experiments or PEER ^60^ factors; 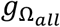 is an n × 1 vector of the total genetic effects of the individuals with 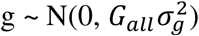 where *G*_*all*_ is the GRM built by all the variants; *W* is the design matrix of variant genotypes (0, 1, 2) and *v* is the variant additive effect; e is the error term. We then saved the eQTL mapping results in the BESD format (https://yanglab.westlake.edu.cn/software/smr/#BESDformat), which is the required data format for SMR. We selected eQTL or sQTL with p < 5e-6 for SMR analysis and a multi-SNP-based SMR test was chosen. Because the RNA-seq data was based on worldwide cattle breeds, we used the 1000-bull whole-genome sequence run7 data ^61^ as the reference panel for SMR.

### Multi-trait meta-analysis of SMR and comparison with MTAO

For a gene or an intron, SMR estimates beta and standard error for a trait, based on the top e/sQTL. Therefore, we can apply equation 21 to 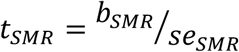 (20) obtained for different traits to test the hypothesis that a gene or an intron has causal effects on more than 1 trait. This meta-analysis also matched the framework of MTAO described above. After calculating the chi-square p-value of multi-trait SMR for each gene and intron similar to equation 18, in each tissue, we count the four following numbers: 1) total number of genes or introns testable between MTAO and SMR; 2) number of genes or introns with p < 0.05 in MTAO; 3) the number of genes or introns with p < 0.05 in multi-trait SMR and 4) the number of genes or introns with p < 0.05 in both MTAO and multi-trait SMR. Then, for each tissue, we used these four counts to generate a contingency table for a Fisher’s exact test [fisher.test(…, alternative=‘greater’) in R v4.0.0] of the significance of the overlap between MTAO and SMR more than expected by random chance. The odds ratio of overlap was obtained from each fisher’s exact test and the p-value was adjusted for multi-testing.

### dN/dS

We retrieved the dN and dS values precalculated by Ensembl (version 99) using R library biomaRt(). dN and dS values were retrieved between cattle (Bos taurus) and humans (Ensembl short label: hsapiens), between cattle and mouse (mmusculus), between cattle and pigs (sscrofa), between cattle and sheep (oaries), between cattle and goat (chircus) and between Bos taurus and Bos indicus (bihybrid, i.e., UOA_Brahman_1). Then the ratio was calculated as dN/dS for all genes participating in the analysis. Only genes with the orthology type as 1-to-1 homology were used in the analysis. The significance of the difference in means of log_2_(dN/dS) between all genes and MTAO prioritised genes was tested in a t-test.

### Relevant tissues for traits

For results from GSOR for each tissue and trait, there is a beta and se, and therefore, *t*_*GSOR*_, estimated for each gene or splicing event. Also, there are results from SMR described above for each gene/splicing which can be combined with results from GSOR to prioritise informative tissues. The squared t-value of a SNP from GWAS can be used to estimate the amount of phenotypic variance explained ^20^. In the current study, to link different tissues to traits, we calculated the following heuristic index: 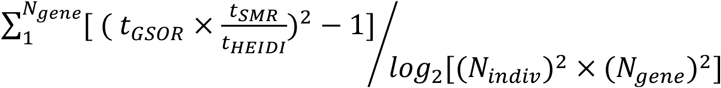 (21) where the magnitude of effects of each gene or intron (|*t*_*GSOR*_|) was adjusted by the magnitude of effect of the SMR test (|*t*_*SMR*_|) and the HEIDI test (|*t*_*HEIDI*_|), so that it is positively related to the causal effects from SMR and negatively related to the LD confounding from HEIDI. The sum of squares was also adjusted for the number of genes and individuals analysed for each tissue and the log scale adjustment made the denominator a linear variable like the numerator. Equation 21 was then used to prioritise informative tissues. Genes and introns were excluded from the analysis if their nominal p-value was > 0.05 in GSOR and SMR and p-value <0.05 in HEIDI test.

## Data and code availability

The newly generated RNA-seq data (356 blood and 268 milk cells) will be made public via NCBI SRA (accession available upon manuscript publication). Other RNA-seq data can be accessed via the CattleGTEx consortium (http://cgtex.roslin.ed.ac.uk/). Summary statistics for genes and splicing events associated with 37 traits of 110,000 cows across 16 tissues are publically available at https://figshare.com/s/c10ffab5abf329b1318f. The DNA sequence data as part of the 1000 Bull Genomes Consortium^61,62^ are available to consortium members and the membership is open. Sequence data of 1832 samples from the 1000 Bull Genome Project have been made publicly available at https://www.ebi.ac.uk/eva/?eva-study=PRJEB42783. DataGene Australia (http://www.datagene.com.au/) are custodians of the raw phenotype and genotype data of Australian farm animals. Access to these data for research requires permission from DataGene under a Data Use Agreement. Other supporting data are shown in the Supplementary Materials of the manuscript.

Code and tutorials for GSOR and MTAO are available at https://github.com/rxiangr/GSOR-and-MTAO. The linear mixed model analysis used GCTA ^56^.

## Acknowledgments

Australian Research Council’s Discovery Projects (DP160101056 and DP200100499) supported R.X., M.E.G. and N.R.W. DairyBio, a joint venture project between Agriculture Victoria (Melbourne, Australia), Dairy Australia (Melbourne, Australia) and the Gardiner Foundation (Melbourne, Australia), funded computing resources used in the analysis. N.R.W. acknowledged funding from National Health and Medical Research Council (NHMRC 1113400 and 1078901). L.F. received funding from European Union’s Horizon 2020 research and innovation programme under the Marie Skłodowska–Curie grant (agreement No. 801215). G.E.L. is partially supported by USDA NIFA AFRI grant numbers 2019–67015-29321 and 2021–67015-33409. A. T. acknowledged funding from the BBSRC through programme grants BBS/E/D/10002070 and BBS/E/D/30002275, MRC research grant MR/P015514/1, and HDR-UK award HDR-9004. The authors also thank the University of Melbourne, Australia for supporting this research. No funding bodies participated in the design of the study nor analysis, interpretation of data, or writing the manuscript. The 1000 Bull Genomes consortium provided access to cattle sequence data and the CattleGTEx consortium provided access to expression data. We thank Professor Naomi R. Wray for a critical read of the manuscript. For the purpose of open access, the authors have applied a CC BY public copyright license to any Author Accepted Manuscript version arising from this submission.

## Author contributions

M.E.G. and R.X. conceived the study. R.X. carried out the main analyses with assistance from M.E.G.. L.F., S.L., Y.G., G.E.L and A.T. assisted in the analysis of data from CattleGTEx. B.A.M. and A.J.C. generated and assisted in the analysis of the new RNA-seq data. R.X. and M.E.G. wrote the paper. R.X., M.E.G., L.F., G.E.L, A.J.C. revised the paper. All authors read and approved the final manuscript.

