## Supplementary Information for "Genetic score omics regression and multi-trait meta-analysis detect widespread *cis*-regulatory effects shaping bovine complex traits"

This file contains:

Supplementary Figures 1-5.

Supplementary Tables 1-6.

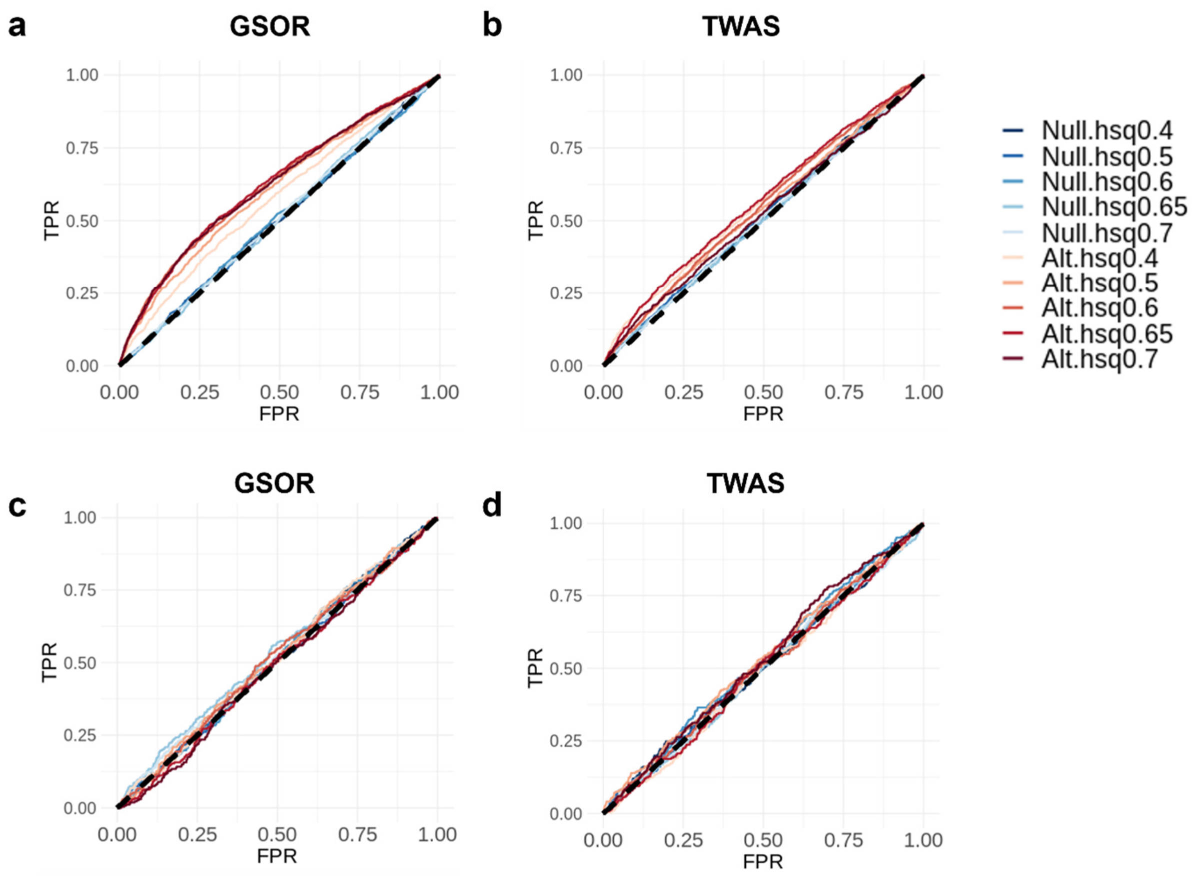


**Supplementary Figure 1**. Comparison of results between GSOR and TWAS using simulated data. Receiver Operating Characteristic (ROC) analysis of results from GSOR and TWAS based on cis+trans predicted values are shown in (**a**) and (**b**), respectively. ROC analysis of results based on trans predicted values are shown in (**c**) and (**d**). 10 scenarios were simulated with varying heritability (hsq) of traits. 5 traits were simulated under the null (Null) where no causal eQTL overlapped with causal QTL and another 5 traits were simulated under the alternative (Alt) scenarios where causal eQTL overlapped with causal QTL for more than 1000 genes.

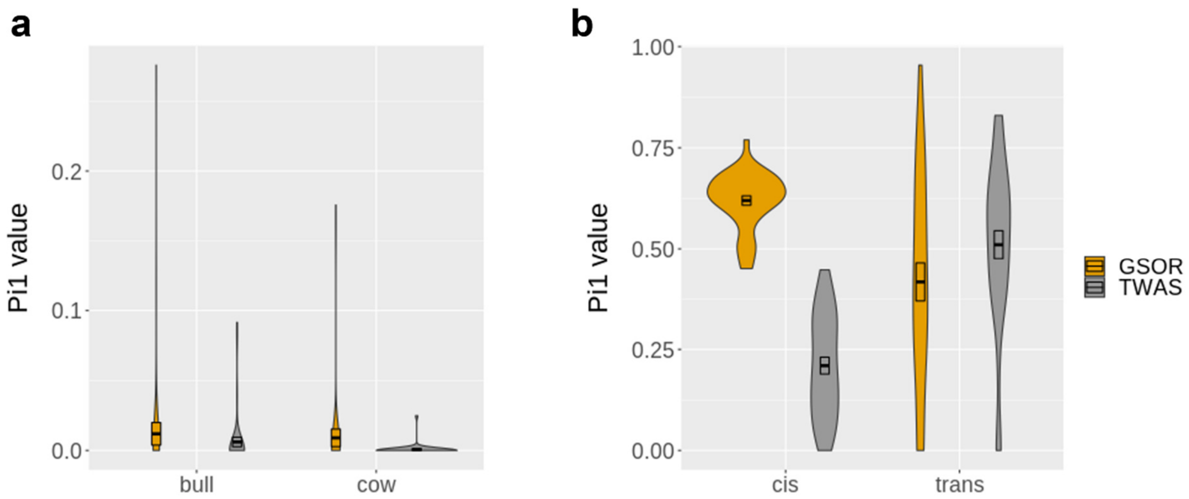


**Supplementary Figure 2**. Comparison of results between GSOR and TWAS using real data. **a**: $\pi_1$ value, an indication of the amount of replicated associations, of GSOR and TWAS across 37 traits between analyses using cis predicted and trans predicted values. Such replication was done in bulls and cows. **b**: $\pi_1$ value of GSOR and TWAS between the two sexes based on cis and trans predicted values.

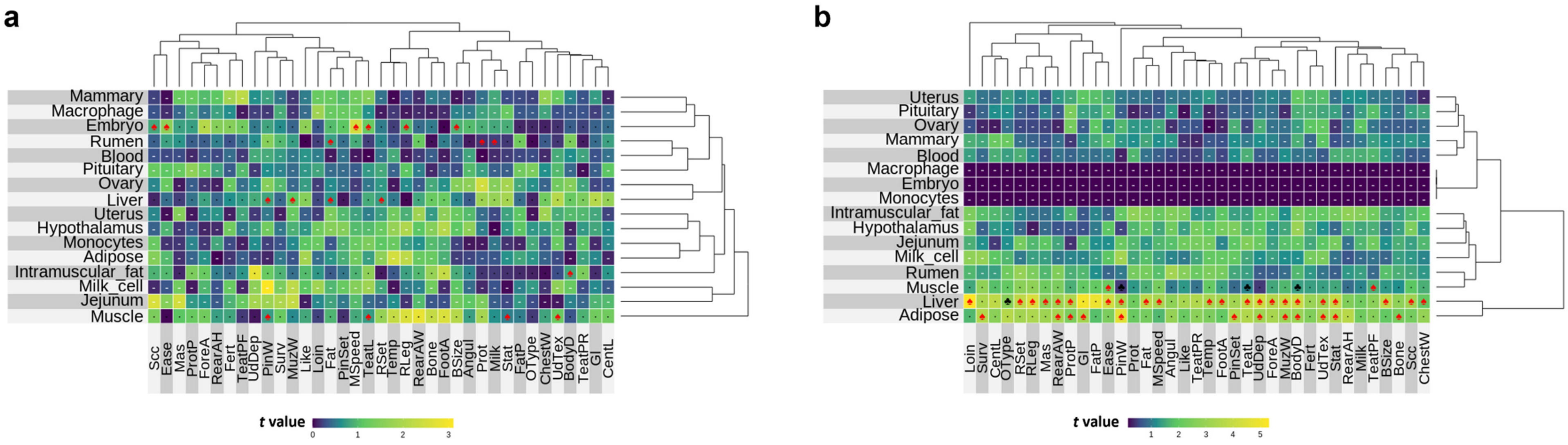


**Supplementary Figure 3**. The heat map of effects of IGF2 expression (**a**) and splicing (**b**) across tissues and traits based on GSOR. In these heat maps, red spades indicate causal effects interred using summary-based Mendelian randomisation (SMR) independent of LD; black hearts indicate the causal effects confounded by LD while black clubs indicate causal effects without testing LD due to not enough SNPs. Black dots represent insignificant SMR test and white hyphens indicate no e/sQTL or QTL can be used for SMR test. The dendrogram represents the hierarchical clustering of effects. The color scale of heatmaps is based on the magnitude of t (be/se) value of GSOR.

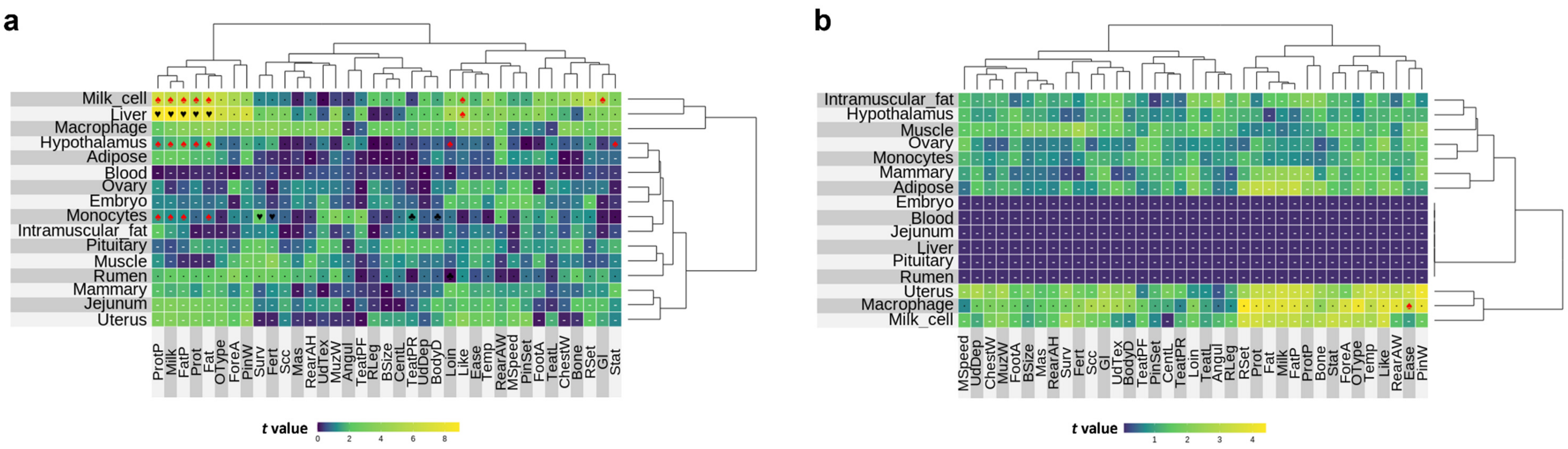


**Supplementary Figure 4**. The heat map of effects of *MGST1* expression (**a**) and splicing (**b**) across tissues and traits based on GSOR. In these heat maps, red spades indicate causal effects interred using summary-based Mendelian randomisation (SMR) independent of LD; black hearts indicate the causal effects confounded by LD while black clubs indicate causal effects without testing LD due to not enough SNPs. Black dots represent insignificant SMR test and white hyphens indicate no e/sQTL or QTL can be used for SMR test. The dendrogram represents the hierarchical clustering of effects. The color scale of heatmaps is based on the magnitude of t (be/se) value of GSOR.

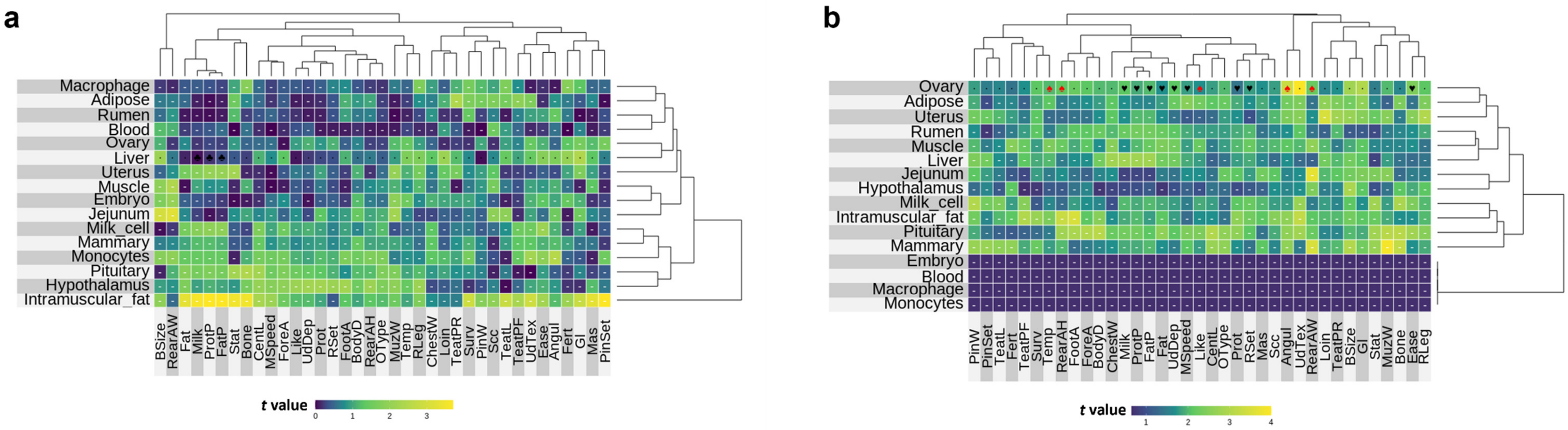


**Supplementary Figure 5**. The heat map of effects of *GHR* expression (**a**) and splicing (**b**) across tissues and traits based on GSOR. In these heat maps, red spades indicate causal effects interred using summary-based Mendelian randomisation (SMR) independent of LD; black hearts indicate the causal effects confounded by LD while black clubs indicate causal effects without testing LD due to not enough SNPs. Black dots represent insignificant SMR test and white hyphens indicate no e/sQTL or QTL can be used for SMR test. The dendrogram represents the hierarchical clustering of effects. The color scale of heatmaps is based on the magnitude of t (be/se) value of GSOR.

**Supplementary Table 1**. cattle traits analysed in the study.cow.N: number of cows for each trait. bull.N: number of bulls for each trait.

| trait full name | short.name | trait.order | cow.N | bull.N |
|---|---|---|---|---|
| protein yield | Prot | tr01 | 76659 | 8097 |
| fat yield | Fat | tr02 | 76659 | 8097 |
| milk yield | Milk | tr03 | 76659 | 8097 |
| protein percentage | ProtP | tr04 | 76659 | 8097 |
| fat percentage | FatP | tr05 | 76659 | 8097 |
| mastitis | Mas | tr06 | 77642 | 8103 |
| somatic cell count | Scc | tr07 | 75429 | 8083 |
| survival | Surv | tr08 | 61056 | 7147 |
| fertility | Fert | tr09 | 56840 | 7254 |
| ease (of birth) | Ease | tr10 | 43970 | 7835 |
| birth size | BSize | tr11 | 43755 | 7827 |
| gestation length | Gl | tr12 | 37214 | 7181 |
| temperament | Temp | tr13 | 37355 | 6966 |
| milking speed | MSpeed | tr14 | 37303 | 6966 |
| likeability | Like | tr15 | 37323 | 6966 |
| stature | Stat | tr16 | 45056 | 7022 |
| chest width | ChestW | tr17 | 45056 | 7022 |
| angularity | Angul | tr18 | 45056 | 7022 |
| bone quality | Bone | tr19 | 45056 | 7022 |
| rear legs set | RSet | tr20 | 45056 | 7022 |
| fore attachment | ForeA | tr21 | 45056 | 7022 |
| rear attachment height | RearAH | tr22 | 45056 | 7022 |
| front teat placement | TeatPF | tr23 | 45056 | 7022 |
| loin strength | Loin | tr24 | 45056 | 7022 |
| udder texture | UdTex | tr25 | 45056 | 7022 |
| central ligament | CentL | tr26 | 45055 | 7022 |
| pin width | PinW | tr27 | 45055 | 7022 |
| foot angle | FootA | tr28 | 45055 | 7022 |
| udder depth | UdDep | tr29 | 45055 | 7022 |
| pin set | PinSet | tr30 | 45054 | 7022 |
| rear attachment width | RearAW | tr31 | 45054 | 7022 |
| muzzle width | MuzW | tr32 | 45054 | 7022 |
| teat length | TeatL | tr33 | 45053 | 7022 |
| body depth | BodyD | tr34 | 45052 | 7022 |
| overall type | OType | tr35 | 44930 | 7022 |
| rear teat placement | TeatPR | tr36 | 44706 | 7022 |
| rear leg view | RLeg | tr37 | 44685 | 7022 |

**Supplementary Table 2**. The number of independent individuals per tissue used for eQTL and sQTL mapping.

| Tissue | N of individuals | Generation |
|---|---|---|
| Blood | 945 | AVR (356) + CattleGTEx (589) |
| Muscle | 699 | CattleGTEx |
| Liver | 576 | CattleGTEx |
| Uterus | 359 | CattleGTEx |
| Macrophage | 295 | CattleGTEx |
| Embryo | 281 | CattleGTEx |
| Milk_cell | 268 | AVR |
| Rumen | 202 | CattleGTEx |
| Mammary | 175 | CattleGTEx |
| Muscle (Cesar et al.) | 171 | CattleGTEx |
| Adipose | 151 | CattleGTEx |
| Ovary | 139 | CattleGTEx |
| Pituitary | 134 | CattleGTEx |
| Monocytes | 113 | CattleGTEx |
| Hypothalamus | 112 | CattleGTEx |
| Jejunum | 105 | CattleGTEx |
| Average | 295 | |
| Total | 4725 | |

AVR: dataset generated by Agriculture Victoria, Australia.
CattleGTEx: data from the Cattle GTEx consortium (https://www.biorxiv.org/content/10.1101/2020.12.01.406280v2).

**Supplementary Table 3**. Summary of results of MTAO. tot.Ngene: total number of genes/intron analysed for each tissue. n.sig.Ngene: number of genes with nominal p-value of Pn (number of traits affected) < 0.05; ave.Pn: average value of Pn. pn.sig.padj.Ngene: number of genes multi-testing adjusted p-value of Pn < 0.05; ave.Pn.padj: average value of Pn using adjusted p-value cutoff. pm.sig.Ngene: number of genes with nominal p-value of Pm (magnitude of multi-trait effects) < 0.05; ave.Pm: average value of Pm. pm.sig.padj.Ngene: number of genes with multi-testing adjusted p-value of Pm < 0.05; ave.Pm.padj: average value of Pm using adjusted p-value cutoff.

| genetype | Tissue | tot.Ngene | pn.sig.Ngene | ave.Pn | pn.sig.padj.Ngene | ave.Pn.padj | pm.sig.Ngene | ave.Pm | pm.sig.padj.Ngene | ave.Pm.padj |
|---|---|---|---|---|---|---|---|---|---|---|
| gene | Adipose | 17660 | 3572 | 8.92 | 2769 | 10.06 | 5003 | 9.30 | 3856 | 9.85 |
| | Blood | 16564 | 2853 | 12.82 | 2554 | 13.74 | 3469 | 13.35 | 3070 | 14.12 |
| | Embryo | 16211 | 1992 | 8.27 | 1154 | 10.34 | 2950 | 8.89 | 1986 | 9.55 |
| | Hypothalamus | 18628 | 3683 | 8.70 | 2854 | 9.78 | 5191 | 9.18 | 3962 | 9.71 |
| | Intramuscular_fat | 16199 | 4170 | 9.81 | 3369 | 10.95 | 5574 | 9.76 | 4690 | 10.21 |
| | Jejunum | 18897 | 4003 | 9.15 | 3136 | 10.30 | 5556 | 9.43 | 4353 | 9.97 |
| | Liver | 16532 | 5629 | 11.18 | 4801 | 12.24 | 7009 | 11.01 | 6275 | 11.43 |
| | Macrophage | 14770 | 2889 | 8.47 | 2211 | 9.53 | 4110 | 9.05 | 3098 | 9.57 |
| | Mammary | 17966 | 3177 | 9.28 | 2523 | 10.39 | 4479 | 9.46 | 3385 | 10.10 |
| | Milk_cell | 16286 | 4746 | 10.04 | 3907 | 11.12 | 6177 | 10.08 | 5350 | 10.49 |
| | Monocytes | 16141 | 1504 | 9.40 | 965 | 11.64 | 2064 | 9.70 | 1469 | 10.56 |
| | Muscle | 15856 | 4435 | 10.36 | 3678 | 11.46 | 5731 | 10.33 | 4917 | 10.81 |
| | Ovary | 18180 | 3054 | 8.52 | 2377 | 9.53 | 4449 | 9.04 | 3266 | 9.60 |
| | Pituitary | 18862 | 3721 | 8.68 | 2824 | 9.84 | 5235 | 9.22 | 4026 | 9.74 |
| | Rumen | 17490 | 3793 | 9.45 | 2973 | 10.67 | 5140 | 9.63 | 4118 | 10.17 |
| | Uterus | 18589 | 4567 | 9.79 | 3696 | 10.91 | 6109 | 9.82 | 5091 | 10.30 |
| intron | Adipose | 214602 | 29800 | 8.05 | 15925 | 10.34 | 45114 | 8.82 | 30074 | 9.48 |
| | Blood | 171580 | 13405 | 11.23 | 9660 | 13.48 | 17497 | 11.61 | 13301 | 12.88 |
| | Embryo | 30132 | 3140 | 7.67 | 1595 | 9.86 | 4944 | 8.63 | 3022 | 9.31 |
| | Hypothalamus | 192856 | 27340 | 7.76 | 14476 | 9.84 | 41671 | 8.69 | 27841 | 9.28 |
| | Intramuscular_fat | 125055 | 16685 | 8.39 | 9485 | 10.64 | 25506 | 8.92 | 17466 | 9.57 |
| | Jejunum | 165578 | 22885 | 8.05 | 12483 | 10.23 | 34553 | 8.81 | 23262 | 9.44 |
| | Liver | 196601 | 34262 | 8.92 | 26276 | 10.11 | 49214 | 9.33 | 36280 | 9.99 |
| | Macrophage | 153138 | 17920 | 7.71 | 9360 | 9.82 | 27825 | 8.65 | 17471 | 9.31 |
| | Mammary | 220783 | 18661 | 8.12 | 9992 | 10.47 | 29552 | 8.81 | 17108 | 9.70 |

| | | | | | | | | | |
|---|---|---|---|---|---|---|---|---|---|
| Milk_cell | 221343 | 49114 | 9.47 | 38991 | 10.63 | 66768 | 9.72 | 53388 | 10.29 |
| Monocytes | 146270 | 10290 | 7.47 | 5032 | 9.64 | 16666 | 8.55 | 8906 | 9.38 |
| Muscle | 172523 | 29626 | 8.64 | 22382 | 9.82 | 42830 | 9.16 | 31149 | 9.79 |
| Ovary | 177820 | 25057 | 7.86 | 13451 | 9.96 | 37725 | 8.74 | 25602 | 9.33 |
| Pituitary | 206952 | 27829 | 7.79 | 14512 | 9.97 | 42671 | 8.71 | 27705 | 9.35 |
| Rumen | 175348 | 22048 | 8.04 | 11971 | 10.25 | 33958 | 8.79 | 22097 | 9.47 |
| Uterus | 212765 | 27569 | 8.34 | 15260 | 10.70 | 41148 | 8.97 | 27230 | 9.71 |

**Supplementary Table 4**. Overlap of prioritised genes/introns between multi-trait meta-analysis of omics-associations (MTAO) and multi-trait summary data-based Mendelian randomization (SMR). N.total: total number of genes or introns testable between MTAO and SMR. N.mtaosig: Number of genes significant in MTAO. N.smrsig: number of genes significant in multi-trait SMR. N.smrsig.mtaosig: Number of genes significant in both MTAO and multi-trait SMR. odds.ratio: fisher's exact test on a contingency table based on N.smrsig.mtaosig, N.mtaosig-N.smrsig.mtaosig, N.smrsig-N.smrsig.mtaosig and N.total-N.mtaosig-N.smrsig +N.smrsig.mtaosig. p.fe.adj: FDR adjusted p-value of fisher's exact test.

| Omics feature | Tissue | N.total | N.mtaosig | N.smrsig | N.smrsig.mtaosig | odds.ratio | p.fe.adj |
|---|---|---|---|---|---|---|---|
| gene expression | Adipose | 1245 | 642 | 65 | 44 | 2.04 | 3.77E-02 |
| | Blood | 6346 | 2390 | 490 | 276 | 2.28 | 3.36E-17 |
| | Embryo | 2557 | 404 | 110 | 22 | 1.35 | 2.73E-01 |
| | Hypothalamus | 2181 | 534 | 83 | 24 | 1.27 | 2.73E-01 |
| | Intramuscular_fat | 2165 | 1061 | 93 | 59 | 1.85 | 2.67E-02 |
| | Jejunum | 1012 | 542 | 49 | 35 | 2.25 | 4.17E-02 |
| | Liver | 5344 | 2991 | 334 | 251 | 2.50 | 7.68E-13 |
| | Macrophage | 1799 | 449 | 52 | 19 | 1.76 | 1.34E-01 |
| | Mammary | 1286 | 699 | 75 | 69 | 10.59 | 4.13E-12 |
| | Milk_cell | 1524 | 1405 | 133 | 130 | 3.94 | 3.77E-02 |
| | Monocytes | 4866 | 593 | 105 | 28 | 2.70 | 3.50E-04 |
| | Muscle | 4597 | 2095 | 259 | 170 | 2.39 | 2.34E-10 |
| | Ovary | 1986 | 361 | 78 | 21 | 1.70 | 1.34E-01 |
| | Pituitary | 3481 | 1019 | 183 | 66 | 1.39 | 1.24E-01 |
| | Rumen | 2937 | 1022 | 134 | 77 | 2.66 | 4.12E-07 |
| | Uterus | 2987 | 1494 | 147 | 103 | 2.44 | 3.81E-06 |
| RNA splicing | Adipose | 9600 | 2377 | 430 | 173 | 2.13 | 3.77E-12 |
| | Blood | 21453 | 6785 | 1571 | 795 | 2.38 | 3.34E-58 |
| | Embryo | 2030 | 243 | 60 | 17 | 3.05 | 1.20E-03 |
| | Hypothalamus | 9505 | 1450 | 340 | 73 | 1.55 | 2.34E-03 |
| | Intramuscular_fat | 11656 | 2396 | 489 | 157 | 1.89 | 4.33E-09 |
| | Jejunum | 7259 | 1372 | 278 | 95 | 2.32 | 4.33E-09 |

| | | | | | | |
|---|---|---|---|---|---|---|
| Liver | 19824 | 6319 | 986 | 434 | 1.73 | 2.15E-15 |
| Macrophage | 14543 | 1899 | 571 | 96 | 1.36 | 4.99E-03 |
| Mammary | 10870 | 2074 | 336 | 142 | 3.26 | 3.45E-22 |
| Milk_cell | 8955 | 7786 | 798 | 747 | 2.33 | 1.49E-09 |
| Monocytes | 9571 | 751 | 296 | 42 | 2.00 | 5.75E-04 |
| Muscle | 18118 | 5108 | 923 | 383 | 1.87 | 4.95E-18 |
| Ovary | 14326 | 2159 | 631 | 129 | 1.48 | 5.75E-04 |
| Pituitary | 15620 | 2644 | 609 | 142 | 1.52 | 1.39E-04 |
| Rumen | 13010 | 2199 | 459 | 133 | 2.07 | 4.15E-10 |
| Uterus | 14658 | 3778 | 602 | 296 | 2.94 | 3.56E-35 |

**Supplementary Table 5**. correlation between tissue ranking and sample size. rho: spearman correlation coefficient; p: raw p-value of rho; p.adjust: FDR adjusted p.value.

| type | tr | rho | p | p.adj |
|---|---|---|---|---|
| gene | Prot | 0.188235 | 0.483876 | 0.628522 |
| | Fat | 0.273529 | 0.304279 | 0.50296 |
| | Milk | 0.382353 | 0.14467 | 0.454856 |
| | ProtP | 0.4 | 0.125879 | 0.454856 |
| | FatP | 0.452941 | 0.080004 | 0.426002 |
| | Mas | 0.173529 | 0.519358 | 0.628522 |
| | Scc | 0.15 | 0.57858 | 0.648711 |
| | Surv | 0.344118 | 0.191942 | 0.454856 |
| | Fert | 0.344118 | 0.191942 | 0.454856 |
| | Ease | 0.241176 | 0.366896 | 0.543006 |
| | BSize | 0.491176 | 0.05558 | 0.411294 |
| | Gl | 0.529412 | 0.037277 | 0.379379 |
| | Temp | 0.529412 | 0.037277 | 0.379379 |
| | MSpeed | 0.261765 | 0.326245 | 0.50296 |
| | Like | 0.102941 | 0.704933 | 0.745215 |
| | Stat | 0.45 | 0.082167 | 0.426002 |
| | ChestW | 0.261765 | 0.326245 | 0.50296 |
| | Angul | 0.170588 | 0.526599 | 0.628522 |
| | Bone | 0.329412 | 0.212614 | 0.454856 |
| | RSet | 0.305882 | 0.248661 | 0.484235 |
| | ForeA | 0.117647 | 0.664435 | 0.723061 |
| | RearAH | -0.02059 | 0.943224 | 0.943224 |
| | TeatPF | 0.173529 | 0.519358 | 0.628522 |
| | Loin | 0.061765 | 0.822182 | 0.84502 |
| | UdTex | 0.335294 | 0.204176 | 0.454856 |
| | CentL | 0.267647 | 0.315146 | 0.50296 |
| | PinW | 0.158824 | 0.556032 | 0.642912 |
| | FootA | 0.361765 | 0.168981 | 0.454856 |
| | UdDep | 0.423529 | 0.103622 | 0.426002 |
| | PinSet | 0.323529 | 0.221281 | 0.454856 |
| | RearAW | 0.641176 | 0.008975 | 0.332092 |
| | MuzW | 0.426471 | 0.101056 | 0.426002 |
| | TeatL | 0.282353 | 0.288413 | 0.50296 |
| | BodyD | 0.335294 | 0.204176 | 0.454856 |
| | OType | 0.223529 | 0.403941 | 0.562275 |
| | TeatPR | 0.220588 | 0.410309 | 0.562275 |
| | RLeg | 0.520588 | 0.041014 | 0.379379 |
| intron | Prot | 0.323529 | 0.221281 | 0.340547 |
| | Fat | 0.25 | 0.349132 | 0.430596 |
| | Milk | 0.441176 | 0.088915 | 0.205617 |
| | ProtP | 0.45 | 0.082167 | 0.202678 |
| | FatP | 0.452941 | 0.080004 | 0.202678 |

| | | | |
|---|---|---|---|
| Mas | 0.488235 | 0.057228 | 0.186955 |
| Scc | 0.314706 | 0.234712 | 0.340547 |
| Surv | 0.582353 | 0.02004 | 0.151279 |
| Fert | 0.714706 | 0.002588 | 0.095772 |
| Ease | 0.482353 | 0.060634 | 0.186955 |
| BSize | 0.388235 | 0.1382 | 0.269126 |
| Gl | 0.311765 | 0.239304 | 0.340547 |
| Temp | 0.105882 | 0.696767 | 0.696767 |
| MSpeed | 0.208824 | 0.436322 | 0.461255 |
| Like | 0.338235 | 0.200041 | 0.321805 |
| Stat | 0.194118 | 0.470031 | 0.483088 |
| ChestW | 0.364706 | 0.165346 | 0.304227 |
| Angul | 0.25 | 0.349132 | 0.430596 |
| Bone | 0.660714 | 0.009043 | 0.151279 |
| RSet | 0.35 | 0.184066 | 0.309566 |
| ForeA | 0.491176 | 0.05558 | 0.186955 |
| RearAH | 0.358824 | 0.17267 | 0.304227 |
| TeatPF | 0.408824 | 0.117166 | 0.252679 |
| Loin | 0.273529 | 0.304279 | 0.402083 |
| UdTex | 0.6 | 0.01597 | 0.151279 |
| CentL | 0.244118 | 0.360918 | 0.430773 |
| PinW | 0.402941 | 0.122925 | 0.252679 |
| FootA | 0.461765 | 0.073768 | 0.202678 |
| UdDep | 0.291176 | 0.27307 | 0.374207 |
| PinSet | 0.226471 | 0.397628 | 0.453502 |
| RearAW | 0.547059 | 0.030594 | 0.151279 |
| MuzW | 0.217647 | 0.416731 | 0.453502 |
| TeatL | 0.217647 | 0.416731 | 0.453502 |
| BodyD | 0.541176 | 0.032709 | 0.151279 |
| OType | 0.561765 | 0.025771 | 0.151279 |
| TeatPR | 0.525 | 0.047104 | 0.186955 |
| RLeg | 0.552941 | 0.028588 | 0.151279 |

**Supplementary Table 6**. cis eQTL used to test SMR and HEIDI for DGAT1 and its neighbor genes of ZNF34 and IQANK1. N number of SNPs.

| N of eQTL for DGAT1 | N of eQTL for ZNF34 | N of eQTL for both DGAT1 and ZNF34 | average LD-r between eQTL of DGAT1 and eQTL of ZNF34 |
|---|---|---|---|
| 1554 | 1341 | 463 | 0.707 |
| **N of eQTL for DGAT1** | **N of eQTL for IQANK1** | **N of eQTL for both DGAT1 and IQANK1** | **average LD-r between eQTL of DGAT1 and eQTL of IQANK1** |
| 1554 | 662 | 317 | 0.708 |
